# Integration of visual information in auditory cortex promotes auditory scene analysis through multisensory binding

**DOI:** 10.1101/098798

**Authors:** Huriye Atilgan, Stephen M. Town, Katherine C. Wood, Gareth P. Jones, Ross K. Maddox, Adrian K.C. Lee, Jennifer K. Bizley

## Abstract

How and where in the brain audio-visual signals are bound to create multimodal objects remains unknown. One hypothesis is that temporal coherence between dynamic multisensory signals provides a mechanism for binding stimulus features across sensory modalities. Here we report that when the luminance of a visual stimulus is temporally coherent with the amplitude fluctuations of one sound in a mixture, the representation of that sound is enhanced in auditory cortex. Critically, this enhancement extends to include both binding and non-binding features of the sound. We demonstrate that visual information conveyed from visual cortex, via the phase of the local field potential is combined with auditory information within auditory cortex. These data provide evidence that early cross-sensory binding provides a bottom-up mechanism for the formation of cross-sensory objects and that one role for multisensory binding in auditory cortex is to support auditory scene analysis.

## Introduction

When listening to a sound of interest, we frequently look at the source. However, how auditory and visual information are integrated to form a coherent perceptual object is unknown. The temporal properties of a visual stimulus can be exploited to detect correspondence between auditory and visual streams (Crosse et al., 2015; Denison et al., 2013; Rahne et al., 2008), can bias the perceptual organisation of a sound scene (Brosch et al., 2015), and can enhance or impair listening performance depending on whether the visual stimulus is temporally coherent with a target or distractor sound stream (Maddox et al., 2015). Together, these behavioural results suggest that temporal coherence between auditory and visual stimuli can promote binding of cross-modal features to enable the formation of an auditory-visual (AV) object (Bizley et al., 2016b).

Visual stimuli can both drive and modulate neural activity in primary and non-primary auditory cortex (Bizley et al., 2007a; Chandrasekaran et al., 2013; Ghazanfar et al., 2005; Kayser et al., 2008; Kayser et al., 2010, Perrodin et al., 2015), but the contribution that visual activity in auditory cortex makes to auditory function remains unknown. One possibility is that the integration of cross-sensory information into early sensory cortex provides a bottom-up substrate for the binding of multisensory stimulus features into a single perceptual object (Bizley et al., 2016b). We have recently argued that binding is a distinct form of multisensory integration that underpins perceptual object formation. We hypothesise that binding is associated with a modification of the sensory representation and can be identified by demonstrating a benefit in the behavioural or neural discrimination of a stimulus feature orthogonal to the features that link crossmodal stimuli (Fig. 1a). Therefore, in order to demonstrate binding, an appropriate crossmodal stimulus should elicit not only enhanced neural encoding of the stimulus features that bind auditory and visual streams (the “binding features”), but that there should be enhancement in the representation of *other* stimulus features (“non-binding features” associated with the source (Fig. 1c).

**Figure 1:**
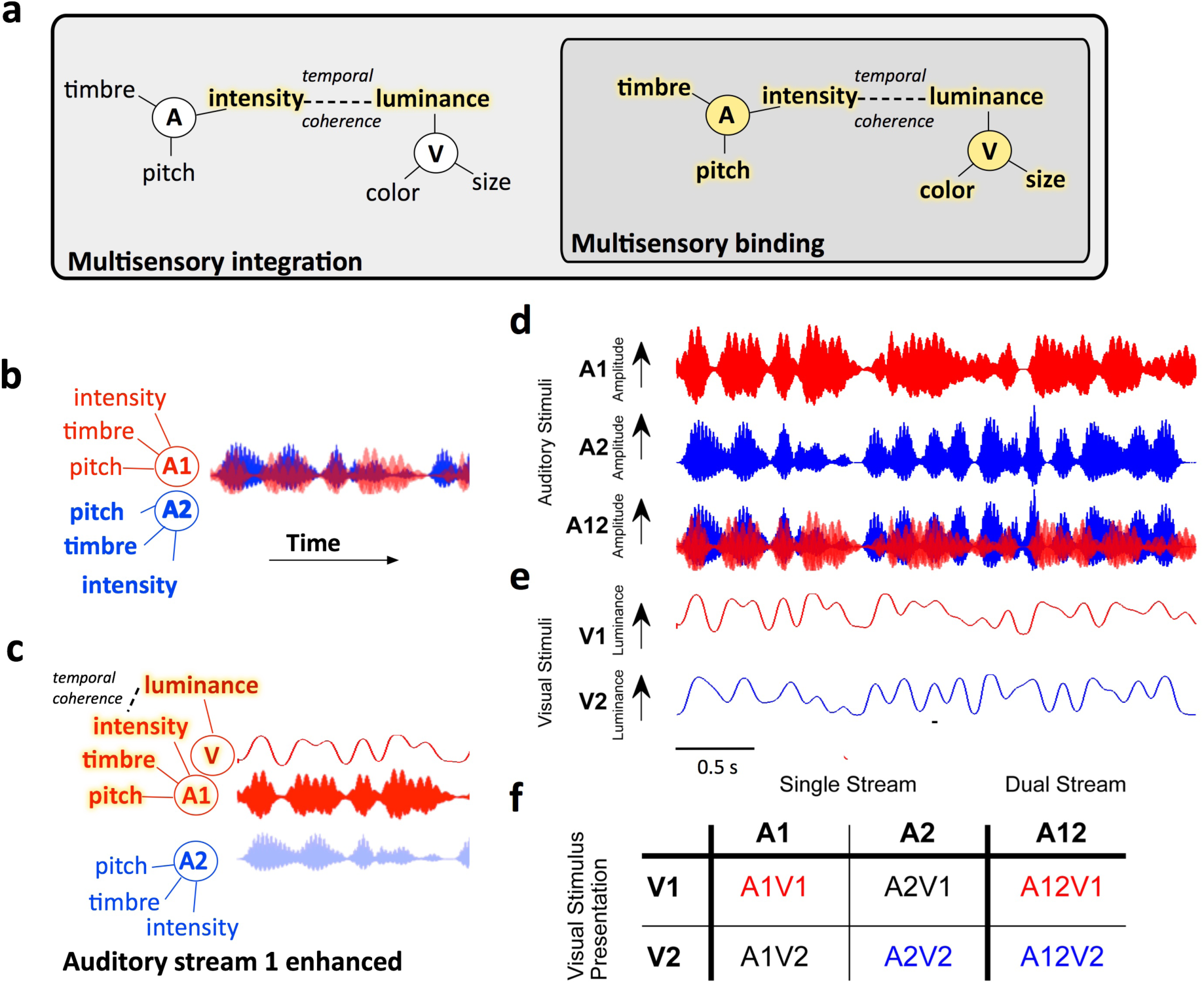
Hypothesis and experimental design. **a** Conceptual model illustrating how binding can be identified as a distinct form of multisensory integration. Multisensory binding is defined as a subset of multisensory integration that results in the formation of a crossmodal object. During binding, all features of the audio-visual object are linked and enhanced - including both those features that bind the stimuli across modalities (here temporal coherence between auditory (A) intensity and visual (V) luminance) and orthogonal features such as auditory pitch and timbre, and visual colour and size. Other forms of multisensory integration would result in enhancement of only the features that promote binding - here auditory intensity and visual luminance. To identify binding therefore requires a demonstration that non-binding features (e.g. here pitch, timbre, colour or size) are enhanced. Enhanced features are highlighted in yellow. **b** When two competing sounds (red and blue waveforms) are presented they can be separated on the basis of their features, but may elicit overlapping neuronal representations in auditory cortex. **c** Hypothesised enhancement in auditory stream segregation when a temporally coherent visual stimulus enables multisensory binding. When the visual stimulus changes coherently with the red sound (A1, top) this sound is enhanced and the two sources are better segregated. Perceptually this would result in more effective auditory scene analysis and an enhancement of the non-binding features. **d** Stimulus design: Auditory stimuli were two artificial vowels (denoted A1 and A2), each with distinct pitch and timbre and independently amplitude modulated with a noisy low pass envelope. **e** Visual stimulus: a luminance modulated white light was presented with one of two temporal envelopes derived from the amplitude modulations of A1 and A2. **f** illustrates the stimulus combinations that were tested experimentally in *single stream* (a single auditory visual pair) and *dual stream* (two sounds and one visual stimulus) conditions. See also supplemental figure 1.

Here we test the hypothesis that the incorporation of visual information into auditory cortex can determine the neuronal representation of an auditory scene through multisensory binding (Fig.1). We demonstrate that when visual luminance changes coherently with the amplitude of one sound in a mixture, auditory cortex is biased towards representing the temporally coherent sound. Consistent with these effects reflecting cross-modal binding, the encoding of sound timbre, a non-binding stimulus feature, is subsequently enhanced in the temporally coherent auditory stream. Finally, we demonstrate that the site of multisensory convergence is in auditory cortex and that visual information is conveyed via the local field potential directly from visual cortex.

## Results

We recorded neuronal responses in the auditory cortex of awake passively listening ferrets (n=9 ferrets, 221 single units, 311 multi-units) in response to naturalistic time-varying auditory and visual stimuli adapted from Maddox et al (2015). The stimuli are designed to share properties with natural speech; they are modulated at approximately syllable rate and, like competing voices, can be separated on the basis of their fundamental frequency (F0, the physical determinant of pitch). These sounds are devoid of any linguistic content permitting the separation of general sensory processing mechanisms from language-specific ones for human listeners. Maddox et al. (2015) used both pure tones and synthetic vowels as stimuli; here we use synthetic vowels as these robustly drive auditory cortical responses in the ferret in neurons with a wide range of characteristic frequencies (Bizley et al., 2009). Ferrets are also well able to distinguish the timbre of artificial vowels (Bizley et al., 2013, Town et al., 2015), and, like human listeners, both ferret behavioural and neural responses show invariant responses to vowel timbre across changes in sound level, location and pitch (Town et al., 2017). We additionally recorded neural responses in medetomidine-ketamine anesthetised ferrets (n=5 ferrets, 426 single units, 772 multi units) which allowed us to entirely eliminate attentional effects and limit the impact of top-down processing. These experiments also permitted longer recording durations for additional control stimuli and enabled simultaneous characterization of neural activity across cortical laminae. In a subset of these animals we were able to reversibly silence visual cortex during recording, in order to determine the origin of visual-stimulus elicited neural changes. Recordings were made in awake freely moving animals while they held their head at a drinking spout but were not engaged in a behavioural task, and allowed us to measure neural activity free from any confounds associated with pharmacological manipulation and in the absence of task-directed attention which would likely engage additional neural circuits.

The stimuli were two auditory streams comprised of two vowels, each with a distinct pitch and timbre (denoted A1: /u/, F0 = 175 Hz and A2: /a/, F0 = 195 Hz, Fig.1) and independently amplitude modulated with a low-pass (<7 Hz) envelope (Fig.1d). A full-field visual stimulus accompanied the auditory stimuli, the luminance of which was temporally modulated with the modulation envelope from one of the two auditory streams (Fig.1e). We tested stimulus conditions in which both auditory streams were presented (“dual stream”) and the visual stimulus was temporally coherent with one or other of the auditory streams (A12V1 or A12V2, Fig.1e). We also tested conditions in which a single AV stimulus pair was presented (‘single stream’ stimuli), where the auditory and visual streams could be temporally coherent (A1V1, A2V2) or independent (A1V2, A2V1), as well as no-visual control conditions.

### Auditory-visual temporal coherence shapes the representation of a sound scene in auditory cortex

We first asked whether the temporal dynamics of a visual stimulus could selectively enhance the representation of one sound in a mixture. We therefore recorded responses to auditory scenes composed of two sounds (A1 and A2), presented simultaneously, with a visual stimulus that was temporally coherent with one or other auditory stream (A12V1 or A12V2). A visual stimulus is known to enhance the representation of the envelope of an attended speech stream in auditory cortex (Zion Golumbic et al., 2013; Park et al., 2016). To test whether we could observe a similar phenomenon in single neurons in the absence of selective attention, we used neural responses to temporally coherent single stream stimuli (i.e. A1V1 and A2V2) to determine to what extent the neural response to the sound mixture was specific to one or other sound stream.

Figure 2 illustrates this approach for a single unit: responses to the temporally coherent single stream AV stimuli (Fig.2a) formed templates which were used to decode the responses to the dual stream stimuli (Fig.2b) using a Euclidean distance based spike pattern classifier. Such an approach is ideally suited for classifying neural responses to time-varying stimuli. Auditory cortical responses to the dual stream stimuli (A12V1 or A12V2) were more commonly decoded as A1V1 when the visual stimulus was V1, and A2V2 when the visual stimulus was V2. Performing this analysis for each neuron in our recorded population yielded similar observations: the coherent auditory stimulus representation was enhanced (Fig.2c,d,f,g) such that auditory cortical responses to dual-stream stimuli most closely resembled responses to the single stream stimulus with the shared visual component.

**Figure 2:**
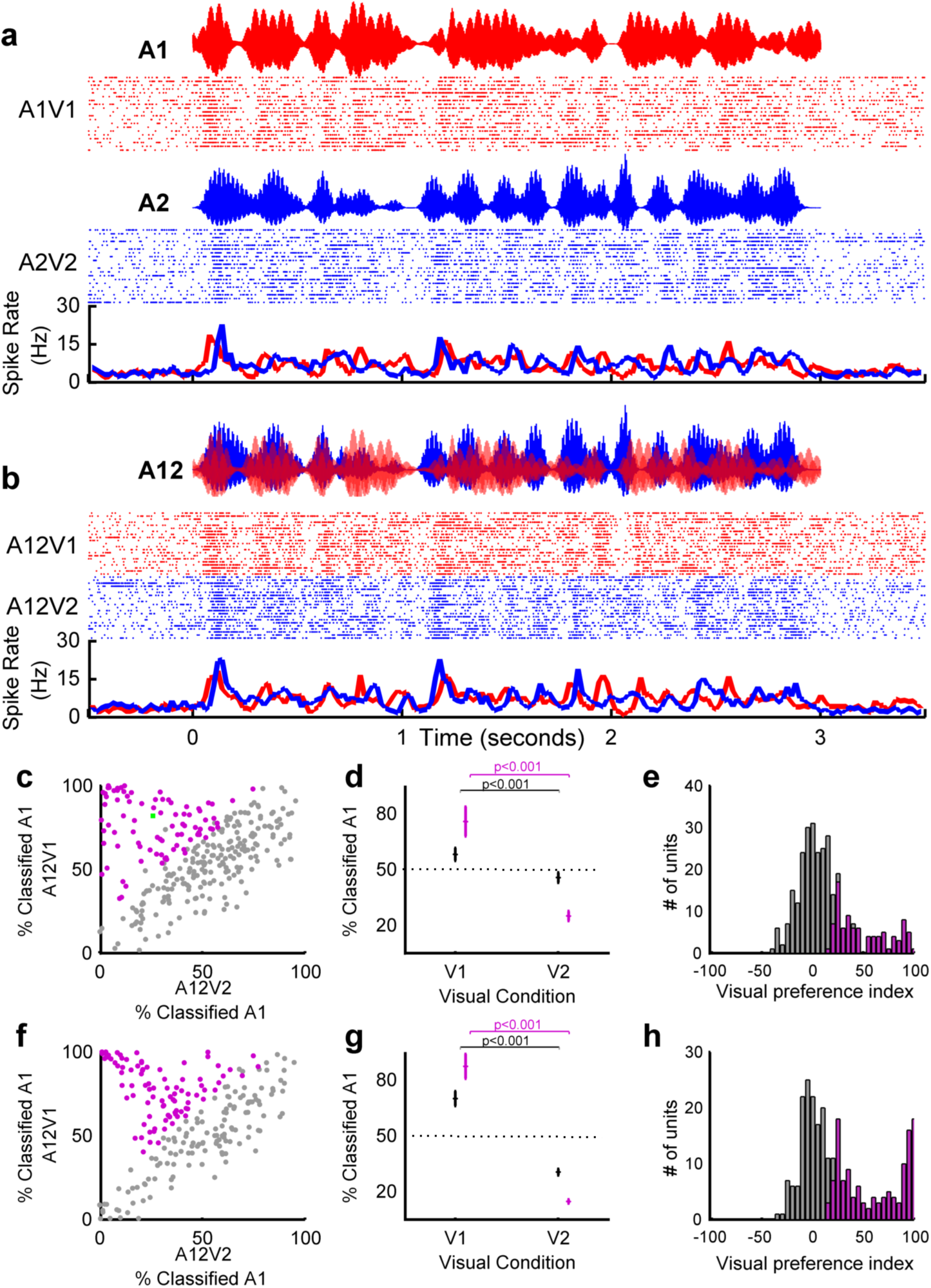
Visual stimuli can determine which sound stream auditory cortical neurons follow in a mixture. Spiking responses from an example unit in response to **a,** single stream AV stimuli used as decoding templates and **b**, dual stream stimuli. In each case rasters and PSTHs are illustrated. When the visual component of the dual stream was V1, the majority of trials were classified as A1V1 (82% (19/23 trials), and A2V2 when the visual stimulus was V2 (26% 6/23, trials) of responses classified as A1V1 (see also green data point in c), yielding a visual preference score of 56%.. **c-h** population data for awake (**c,d,e** 271 units) and anesthetised (**f,g,h** 331 units) datasets. In each case the left panel (**c,f**) shows the distribution of decoding values according to the visual condition, the middle panel (**d,g**) shows the population mean (± SEM) projecting onto the vertical axis of panel c / f for V1 condition, and horizontal axis of panel c / f for the V2 condition. **e,h** shows the visual preference index (VPI). Units in which the VPI was significantly >0 are coloured purple. Pairwise comparisons revealed significant effect of visual condition on decoding in all datasets: Awake: All: t_540_=6.1,p=2.3e-09 (n =271), Sig VPI: t_180_=18.8 p = 2.0e-44 (n=91). Anesthetised: All: t_660_=9.5,p=3.3e-20 (n=331), Sig. VPI: t_348_ = 38.9, p =1.2e-128 (n =175) See also supplemental figures 2-4.

To quantify whether the responses of individual units were significantly influenced by the visual stimulus identity, we first calculated a visual preference index (VPI) as the difference between the percentage of A12V1 trials labelled A1 and the percentage of A12V2 trials labelled A1. Units which were fully influenced by the identity of the visual stimulus would have a visual preference score of 100, while those in which the visual stimulus did not influence the response at all would have a score of 0 (Fig. 2e,h). We assessed the significance of observed VPI scores using a permutation test (p < 0.05).to revealed that 33.6% of driven units recorded in awake animals (91/271 units) and 52.9% of units in the anesthetised dataset (175/331 units) had responses to dual stream stimuli significantly influenced by the visual stimulus.

Modulation of dual stream responses by visual stimulus identity was not simply a consequence of the shared visual component of single stream and dual stream stimuli and was observed in neurons in which visual or auditory identity could be decoded (example response from a unit in which only auditory stimulus identity could be decoded: Fig.S2). If this effect was only apparent in visual neurons in auditory cortex, then eliminating the visual element of the single stream stimuli should impair decoding performance for the dual stream stimuli. Additional control experiments (n=89 driven units, awake animals) demonstrated that this was not the case: the enhancement of the temporally coherent sound in the sound mixture was evident whether dual stream stimuli (A12V1 and A12V2) were decoded using responses to auditory-only stimuli (A1 or A2) or auditory-visual stimuli (A1V1, A2V2 etc.). Within this control data (Example unit: Fig. 3a,b, population data Fig 3c-h), 32 units had a significant VPI scores when dual stream responses were decoded from an auditory-only single stream templates and 31 units when decoded with an auditory-visual template. Furthermore, the distribution of VPI values was statistically indistinguishable for decoding dual stream responses with A-only or AV templates (Kolmogorov–Smirnov test: all units, p= 0.9016; units with visual preference scores significantly >0, p > 0.9998), and the distribution of values in Fi.3d was statistically indistinguishable from that in Fig.2e (p=0.0864). We also determined that removing the visual stimulus from the dual-stream condition eliminated any decoding difference in responses observed (Fig.3h). A two-way repeated measures ANOVA on decoded responses with factors of visual stream (V1, V2, no visual), and template type (AV or A) demonstrated a significant effect of visual stream identity on dual stream decoding (F(2, 528) = 19.320, p < 0.001), but there was no effect of template type (F(1,528) = 0.073, p = 0.787) or interaction between factors (F(2,528) = 0.599, p = 0.550). Post-hoc comparisons revealed that without visual stimulation, there was no tendency to respond preferentially to either stream, but that visual stream identity significantly influenced the classification of dual stream responses.

**Figure 3:**
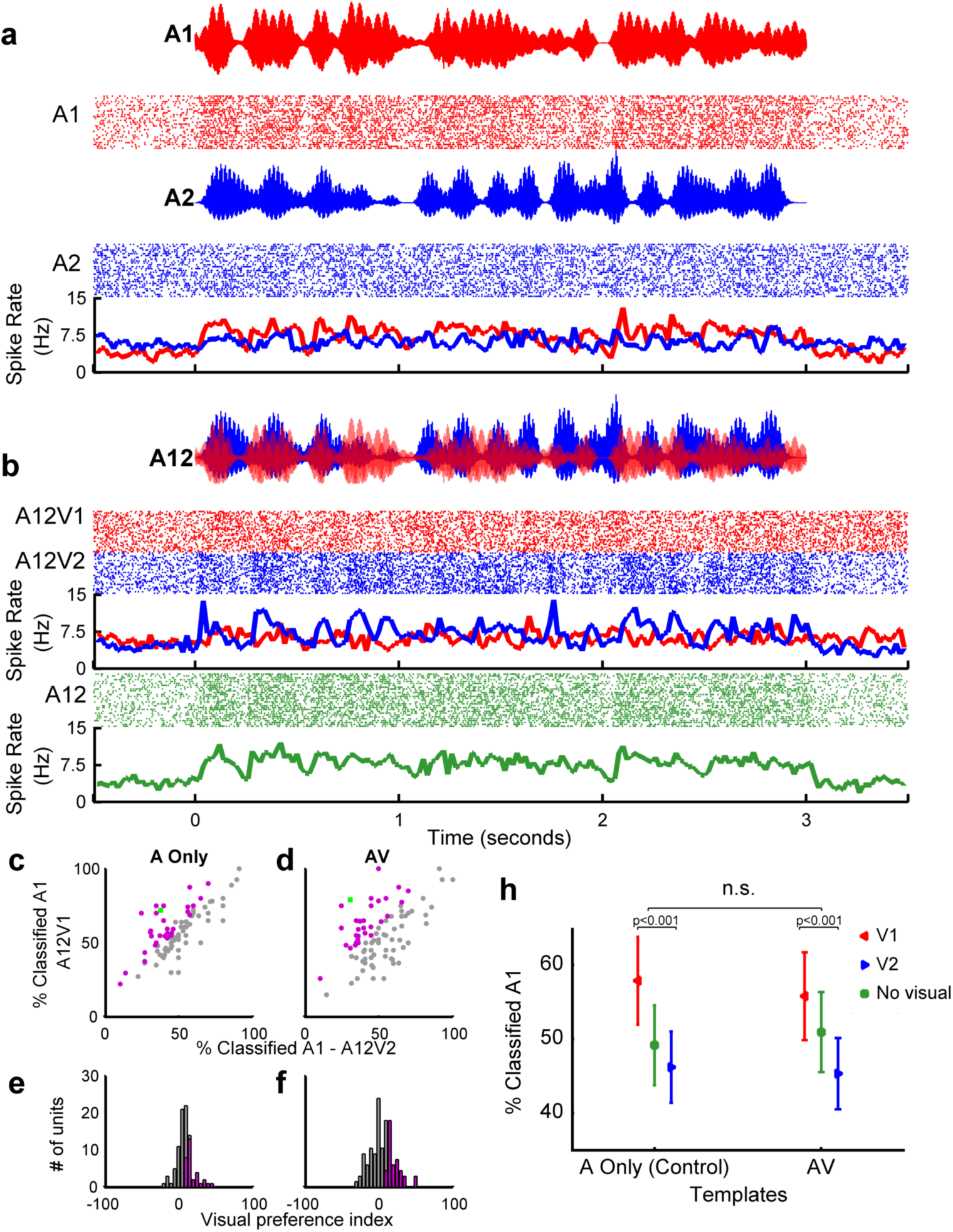
Visual stimuli shape the neural representation of an auditory scene. In an additional control experiment (n=89 units recorded in awake animals), the responses to coherent AV and auditory-only (A Only) single stream stimuli were used as templates to decode dual stream stimuli either accompanied by visual stimuli (V1/V2) or in the absence of visual stimulation (no visual). Spiking responses from an example unit in response to **a,** single stream auditory stimuli which were used as decoding templates to decode the responses to dual stream stimuli in **b**, in each case the auditory waveform, rasters and PSTHs are shown. In this example, when decoded with AV templates: 79% (22/28) of responses were classified as A1 when the visual stimulus was V1, and 32 % of responses (9/28) were classified as A1 when the visual stimulus was V2, yielding a VPI score of 47%. When decoded with A-only templates the values were 75% when V1 (22/28) and 35% when V2 (10/28), yielding a VPI of 40%. For comparison the auditory-only condition (A12) is shown in **c. d**, population data showing the proportion of responses classified as A1 when the visual stimulus was V1 or V2 when decoded with auditory-only templates or auditory visual templates. **e,f,** resulting VPI scores. **h,** Mean (± SEM) values for these units when decoded with A-only templates, AV templates (as in Fig.2) or in the absence of a visual stimulus. The green data point in **d** depicts the example in **a, b**.

Analysis of recording site locations demonstrated that in the awake animals recordings in the Posterior Ectosylvian Gyrus (PEG, which contains two tonotopic secondary fields) were most strongly influenced by the visual stimulus (Fig.S3b). In anesthetised animals the magnitude of the visual preference scores was similar to that of awake animals in the primary fields, but was not significantly different across cortical areas (Fig.S3e). In both awake and anesthetised animals units that were classified as ‘visual-discriminating’ (see Fig.5/methods) and ‘auditory-discriminating’ were influenced by the visual stimulus, with the magnitude of the effects being greatest in the visual-discriminating units. In anesthetised animals we confirmed using noise bursts and light flashes that a substantial proportion of visual-discriminating and auditory-discriminating units were auditory-visual (of 136 visual discriminating units with a significant VPI, 19 were categorised as auditory, 39 as visual and 78 as auditory-visual, of 39 auditory-discriminating units with significant VPI values 21 were auditory, 2 were visual and 16 were auditory visual Fig. S3i). The ability of auditory-visual temporal coherence to enhance one sound in a mixture was observed across all cortical layers (anesthetised dataset; layers defined by current source density analysis, see methods, Fig.S3f), but was strongest in the supra-granular layers (Fig.S3g). Finally, we observed these effects in both single and multi-units (Fig.S5a,b).

### Auditory-visual temporal coherence enhances non-binding sound features

A hallmark of an object-based rather than feature-based representation is that all stimulus features are bound into a unitary perceptual construct, including those features which do not directly mediate binding (Desimone and Duncan, 1995). We predicted that binding across modalities would be promoted via synchronous changes in auditory intensity and visual luminance (Fig.1b, S1) and observed that the temporal dynamics of the visual stimulus enhanced the representation of temporally coherent auditory streams (Fig.2c-h and 3d-f). To determine whether temporal synchrony of visual and auditory stimulus components also enhanced the representation of orthogonal stimulus features and thus fulfil a key prediction of binding (Bizley et al., 2016b), we introduced brief timbre perturbations into our acoustic stimuli (two in each of the A1 and A2 streams). Each deviant lasted for 200 ms during which the spectral timbre smoothly transitioned to the identity of another vowel and back to the original. It is important to note that neither the amplitude of the auditory envelope nor the visual luminance were informative about whether, or when, a change in sound timbre occurred (Fig.S1). Such timbre deviants could be detected by human listeners and were better detected when embedded in an auditory stream that was temporally coherent with an accompanying visual stimulus (Maddox et al., 2015). We hypothesised that a temporally coherent visual stimulus would enhance the representation of timbre deviants in the responses of auditory cortical neurons.

To isolate neural responses to the timbre change from those elicited by the on-going amplitude modulation, we extracted 200 ms epochs of the neuronal response during the timbre deviant and compared these to epochs from stimuli without deviants that were otherwise identical (Fig.S1). We observed that the spiking activity of many units differed between deviant and no-deviant trials (e.g. Fig.4a, Fig.S6) and we were able to discriminate deviant from no-deviant trials with a spike pattern classifier. For each neuron, our classifier reported both the number of deviants that could be detected (i.e. discriminated better than chance as assessed with a permutation test, the maximum is 4, two per auditory stream), and a classification score (where 100% implies perfect discrimination, and 50% chance discrimination, averaged across all deviants for any unit in which at least one deviant was successfully detected). We first considered the influence of temporal coherence between auditory and visual stimuli on the representation of timbre deviants in the single stream condition (A1V1, A1V2 etc.). We found that a greater proportion of units detected at least one deviant when the auditory stream in which deviants occurred was temporally coherent with the visual stimulus relative to the temporally independent condition. This was true both for awake (Fig. 4b; Pearson chi-square statistic, χ^2^ = 322.617, p < 0.001) and anesthetised animals (Fig. 4e; χ^2^ = 288.731, p < 0.001). For units that detected at least one deviant, discrimination scores were significantly higher when accompanied by a temporally coherent visual stimulus (Fig.4c, awake dataset, pairwise t-test t_300_ = 3.599 p < 0.001; Fig. 4f, anesthetised data t_262_ = 4.444 p < 0.001).

**Figure 4:**
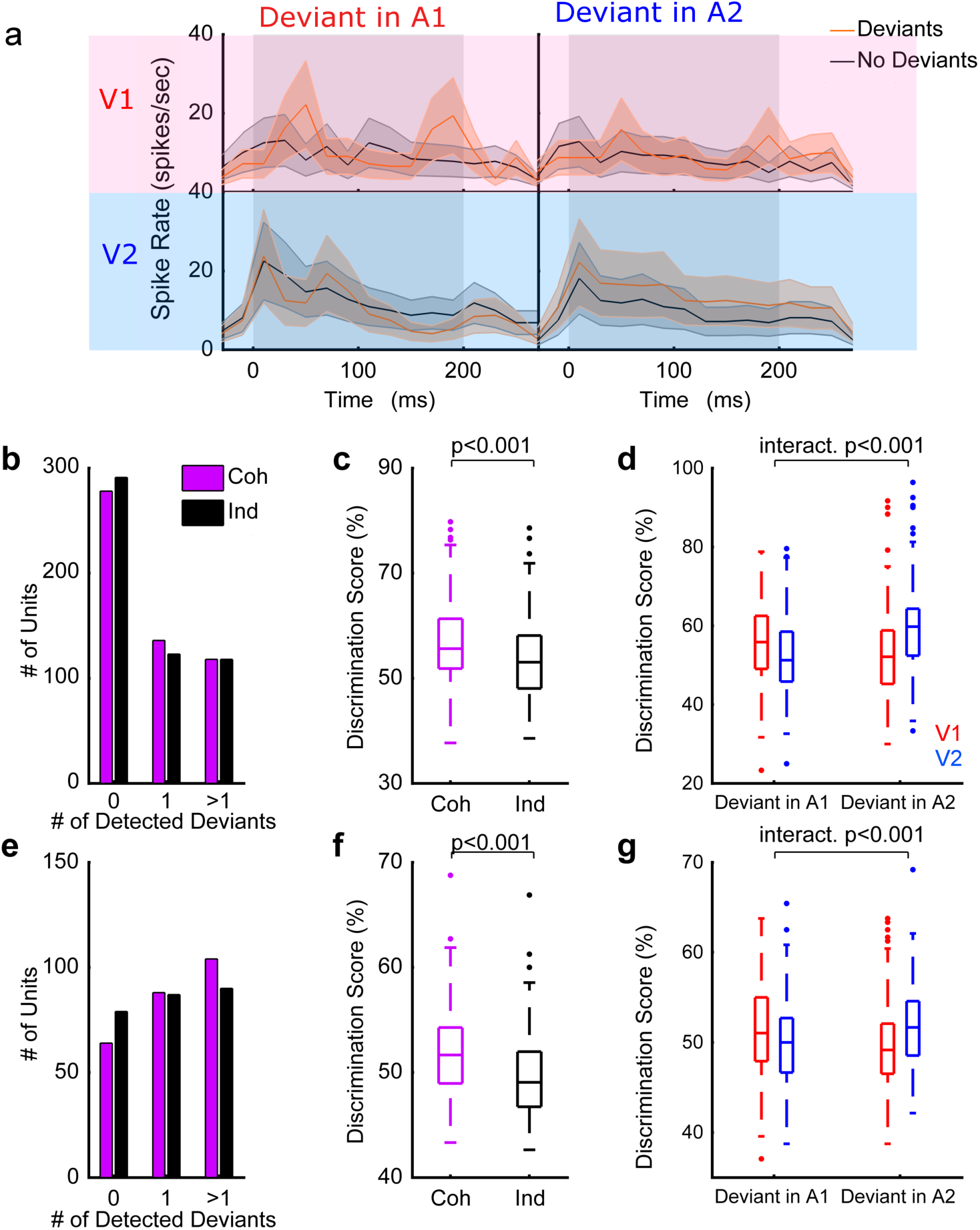
Temporally coherent changes in visual luminance and auditory intensity enhance the representation of auditory timbre. **a** Example unit response (from the awake dataset) showing the influence of visual temporal coherence on spiking responses to dual stream stimuli with (red PSTH) or without (black PSTH) timbre deviants. **b-d** timbre deviant discrimination in the awake dataset. Two deviants were included in each auditory stream giving a possible maximum of 4 per unit **b,** Histogram showing the number of deviants (out of 4) that could be discriminated from spiking responses **c,** Box plots showing the timbre deviant discrimination scores in the single stream condition across different visual conditions (Coh: coherent, ind: independent). The boxes show the upper and lower quartile values, and the horizontal lines indicates the median, the whiskers depict the most extreme data points not considered to be outliers (which are marked as individual symbols). **d,** Discrimination scores for timbre deviant detection in dual stream stimuli. Discrimination scores are plotted according to the auditory stream in which the deviant occurred and the visual stream that accompanied the sound mixture. V1 stimuli are plotted in red, and V2 stimuli in blue; therefore for d and g the boxplots at the far left and right of the plot represent the cases in which the deviants occurred in an auditory stream which was temporally coherent with the visual stimulus while the central two boxplots represent the discrimination of deviants occurring in the auditory stream which was temporally independent of the visual stimulus. **e-g** show the same as **b-d** but for the anesthetised dataset. See also supplemental figure 6.

We also observed an enhancement in the representation of timbre changes in the context of a sound scene (Fig 4d,g): timbre changes were more reliably encoded when the sound stream in which they were embedded was accompanied by a temporally coherent visual stimulus. We performed a two-way repeated measures ANOVA on deviant discrimination performance with visual condition (V1/V2) and the auditory stream in which the deviants occurred (A1/A2) as factors. We anticipated that enhancement of the representation of timbre deviants in the temporally coherent auditory stream would be revealed as a significant interaction term in the dual stream data. Significant interaction terms were seen in both the awake (Fig.4d, F(1,600) = 29.138, p < 0.001) and anesthetised datasets (Fig.4g,F(1,524) = 16.652, p < 0.001). We also observed significant main effects of auditory and visual conditions in awake (main effect of auditory stream, F(1,600) = 4.565, p = 0.033; main effect of visual condition, F(1,600) = 2.650, p = 0.010) but not anesthetised animals (main effect of auditory stream, F(1,524) = 0.004, p = 0.948; main effect of visual condition, F(1,524) =1.355, p = 0.245).

Finally, to determine whether a temporally coherent visual stimulus enhanced the representation of non-binding features relative to auditory-alone stimuli, we collected additional control data (3 animals, 39 driven units) in which single stream stimuli were presented with, or without a temporally coherent visual stimulus. These data (Fig.S6a-c) confirmed that the presence of a visual stimulus enhanced the encoding of timbre deviants relative to the auditory-only condition. The magnitude of the influence of auditory-visual temporal coherence on timbre deviant encoding was equivalent in single and multi units (Fig.S5c,d).

Together these data demonstrate the predicted enhancement in the neural representation of both binding (i.e. auditory amplitude) and non-binding features (here auditory timbre) that are orthogonal to those that promote binding between auditory and visual streams, meaning the effects we observe in auditory cortex fulfil our definition of multisensory binding. Next we turn to the question of how these effects are mediated, and whether they emerge within or outside of auditory cortex.

### Auditory cortical spike patterns differentiate dynamic auditory-visual stimuli more effectively when stimuli are temporally coherent

We used the responses to single stream stimuli to classify neurons according to whether they were dominantly modulated by auditory or visual temporal dynamics. To determine whether the auditory amplitude envelope reliably modulated spiking, we used a spike-pattern classifier to decode the auditory stream identity, collapsed across visual stimulus (i.e. we decoded auditory stream identity from the combined responses to A1V1 and A1V2 stimuli and the combination of A2V1 and A2V2 responses). An identical approach was taken to determine if neuronal responses reliably distinguished visual modulation (i.e. we decoded visual identity from the combined responses to A1V1 and A2V1 stimuli and the combined responses elicited by A1V2 and A2V2). Neuronal responses which were informative about auditory or visual stimulus identity at a level better than chance (estimated with a bootstrap resampling) were classified as auditory-discriminating (Fig. 5a-b) and / or visual-discriminating (Fig. 5c-d) respectively.

**Figure 5:**
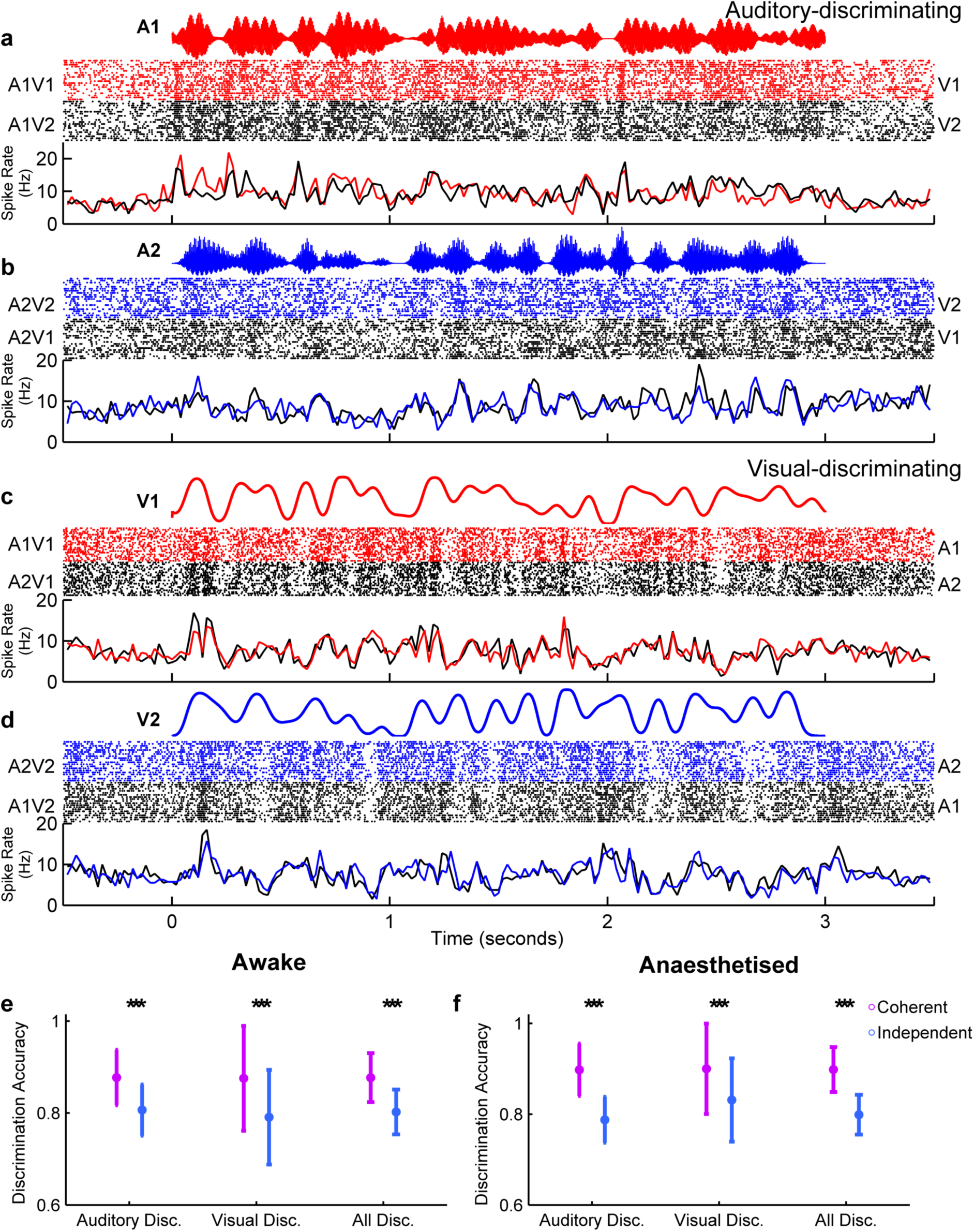
Auditory-visual temporal coherence enhances neural coding in auditory cortex. A pattern classifier was used to determine whether neuronal responses were informative about auditory or visual stimuli. The responses to single stream stimuli are shown for two example units, with responses grouped according to the identity of the auditory (**a, b**, for an auditory discriminating unit) or visual stream (**c, d**, for a visual discriminating unit). In each case the stimulus amplitude (a,b) / luminance (c,d) waveform is shown in the top panel with the resulting raster plots and PSTHs below. **e, f**: Decoder performance (mean ± SEM) for discriminating stimulus identity (coherent: A1V1 vs. A2V2, purple; independent: A1V2 vs. A2V1, blue) in auditory and visual classified units recorded in awake (e) and anaesthetised (f) ferrets. Pairwise comparisons for decoding of coherent versus independent stimuli (*** indicates p < 0.001).

In awake animals, 39.5% (210/532) of driven units were auditory-discriminating, 11.1% (59/532) were visual-discriminating, and only 0.4% (2/532) discriminated both auditory and visual stimuli. Overall a smaller proportion of units represented the identity of auditory or visual streams in the anesthetised dataset: 20.2% (242/1198) were auditory-discriminating, 6.8% (82/1198) were visual discriminating, and 0.6% (7/1198) discriminated both. Using simple noise bursts and light flashes in anesthetised animals revealed that the classification of units as visual / auditory discriminating based on the single stream stimuli selected a subset of light and/or sound driven units and that the proportions of auditory, visual and AV units recorded in our sample were in line with previous studies from ferret auditory cortex (65.1% (328/504) of units were driven by noise bursts, 16.1% (81/504) by light flashes and 14.1% (71/504) by both). When considering the units which were classified as auditory or visual discriminating based on single stream stimuli, and for which we recorded responses to noise bursts and light flashes, 53% (160/307) were classified as auditory, 17% (53/307) as visual and 31% (94/307) as auditory-visual when classified with simple stimuli (see also Fig.S3i).

We hypothesised that the effects we observed in the dual-stream condition might be a consequence of temporal coherence between auditory and visual stimuli enhancing the discriminability of neural responses. We confirmed this prediction by using the same spike pattern decoder to compare our ability to discriminate temporally coherent (A1V1 vs. A2V2) and temporally independent (A1V2 vs. A2V1) stimuli (Fig.5e,f): Temporally coherent AV stimuli produced more discriminable spike patterns than those elicited by temporally independent ones in both awake (Fig. 5e, pairwise t-test, auditory-discriminating t_418_ = 11.872, p < 0.001; visual-discriminating t_116_ = 6.338, p < 0.001; All t_540_ = 13.610, p < 0.001) and anesthetised recordings (Fig.5f, auditory-discriminating t_482_ = 17.754, p < 0.001; visual-discriminating t_162_ = 8.186, p < 0.001; All t_664_ = 19.461, p < 0.001). We further determined that neither the mean nor maximum evoked spike rates were different between trials in response to temporally coherent and temporally independent auditory visual stimuli (Fig. S4). We also observed that the impact of auditory-visual temporal coherence was stronger in single units than multiunits in the awake dataset (Fig.S5e). Therefore the improved discrimination ability observed in response to temporally coherent auditory-visual stimuli is most likely to arise due to an increase in the reliability with which a spiking response occurred.

### Dynamic visual stimuli elicit reliable changes in LFP phase

Temporal coherence between auditory and visual stimulus streams results in more discriminable spike trains in the single stream condition, and an enhancement of the representation of the temporally coherent sound when that sound forms part of an auditory scene. What might underlie the increased discriminability observed for temporally coherent cross-modal stimuli? The phase of on-going oscillations determines the excitability of the surrounding cortical tissue (Azouz and Gray, 1999; Okun et al., 2010; Szymanski et al., 2011). LFP phase is reliably modulated by naturalistic stimulation (Chandrasekaran et al., 2010; Kayser et al., 2009; Luo and Poeppel, 2007b; Ng et al., 2012; Schyns et al., 2011) and has been implicated in multisensory processing (Golumbic et al., 2013; Lakatos et al., 2007). We hypothesised that sub-threshold visual inputs could modulate spiking activity by modifying the phase of the local field potential such that, when visual-stimulus induced changes in LFP phase coincided with auditory-stimulus evoked activity, the spiking precision in auditory cortex was enhanced.

Stimulus-evoked changes in the local field potential (LFP) were evident from the recorded voltage traces, and analysis of inter-trial phase coherence demonstrated that there were reliable changes in phase across repetitions of identical AV stimuli (Fig.6a,b). To isolate the influence of visual activity on the LFP at each recording site, and address the hypothesis that visual stimuli elicited reliable changes in the LFP, we calculated phase and power dissimilarity functions for stimuli with identical auditory signals but differing visual stimuli (Luo and Poeppel, 2007b). Briefly, this analysis assumes that if the phase within a particular frequency band differs systematically between responses to two different stimuli, then inter-trial phase coherence (ITPC) across repetitions of the same stimulus will be greater than across randomly selected stimuli. For each frequency band in the LFP, we therefore compared “within-stimulus” ITPC for responses to each stimulus (A1 stream Fig. 6c; A2 stream Fig. 6d) with “across-stimulus” ITPC calculated from stimuli with *identical auditory components* but randomly selected visual stimuli (e.g. randomly drawn from A1V1 and A1V2). The difference between within-stimulus and across-stimulus ITPC was then calculated across frequency and described as the phase dissimilarity index (PDI) (Fig.6e,f, single site example, k,l population data) with positive PDI values indicating reliable changes in phase coherence elicited by the visual component of the stimulus. Importantly, because both test distributions and the null distribution contain identical sounds any significant PDI value can be attributed directly to the visual component of the stimulus.

**Figure 6:**
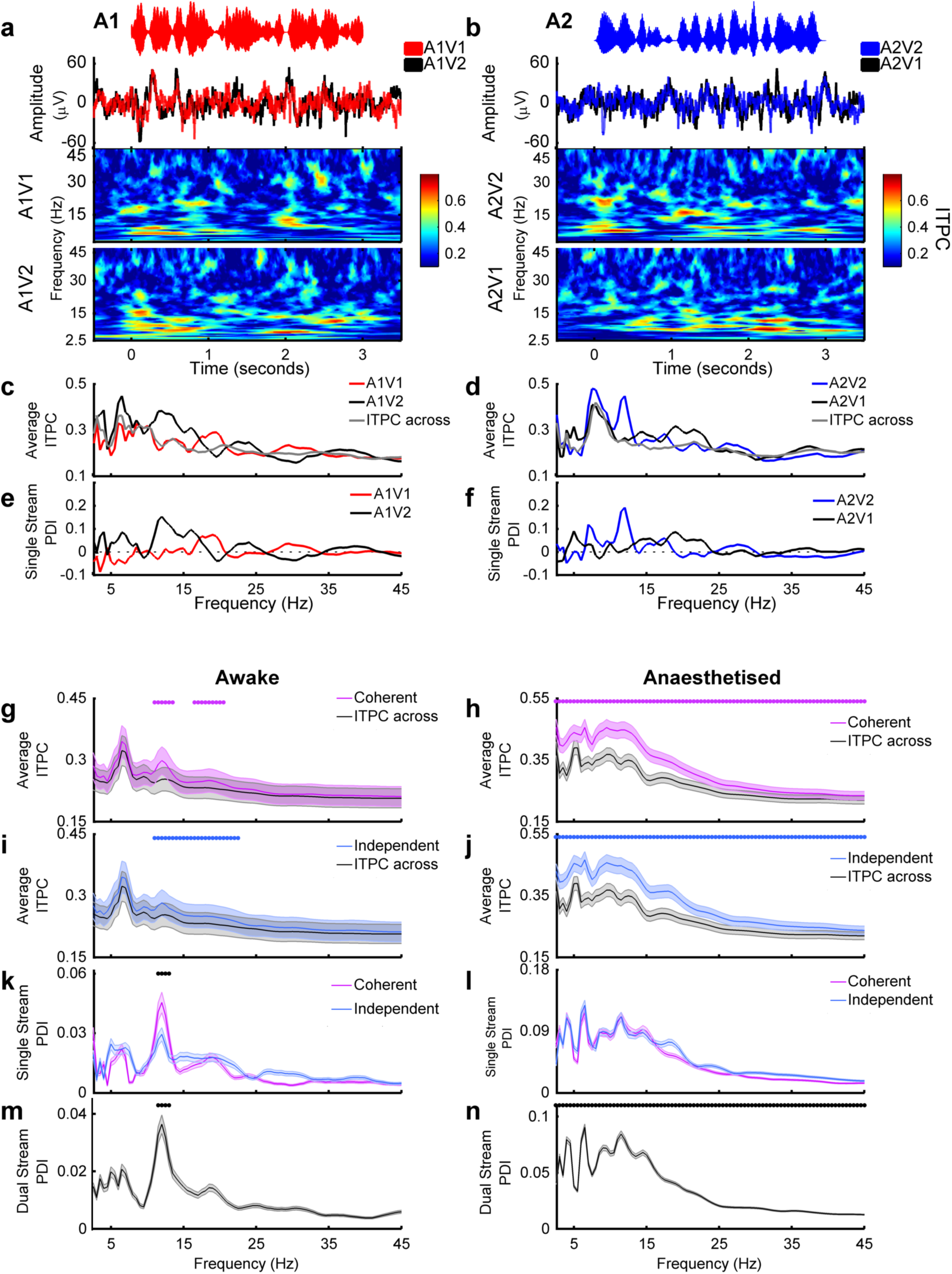
Visual stimuli elicit reliable changes in the phase of the local field potential. **a, b** Example LFP responses to each single stream auditory stimulus across visual conditions. Data obtained from the recording site at which multiunit spiking activity discriminated auditory stream identity in Fig. 5a and b. The amplitude waveforms of the stimuli are shown in the top panel, with the evoked LFP underneath (mean across 21 trials). The resulting inter-trial phase coherence (ITPC) values are shown in the bottom two panels (**c, d**). ITPC was calculated for coherent and independent AV stimuli separately and compared to a null distribution (ITPC across). **e,f** Single stream phase dissimilarity values (PDI) were calculated by comparing ITPC within values to the ITPC across null distribution. **g,I** Population mean inter-trial phase coherence (ITPC) values across frequency for coherent (**g**, significant frequencies 10.5-13, 16-20 Hz) and independent (**i** significant frequencies 10.5-22 Hz) conditions. Dots indicate frequencies at which the ITPC-within values were significantly greater than the ITPC-across values (Pairwise t-test, α = 0.0012, Bonferroni corrected for 43 frequencies). **k:** Mean (±SEM) single stream phase dissimilarity index (PDI) values for coherent and independent stimuli in animals. Black dots indicate frequencies at which the temporally coherent single stream PDI is significantly greater than in the independent conditions (p < 0.001, significant frequencies 10.5-12.5). **h,j,I** as **g,i,k** for anesthetised dataset. **m, n,** mean ±SEM dual stream PDI values for awake (**m**, significant frequencies 10.5-12.5) and anesthetised (n) datasets.

We calculated PDI values for each of the four single stream stimuli and grouped conditions by coherency (coherent: A1V1 / A2V2, or independent: A1V2 / A2V1). To determine at which frequencies the across-trial phase reliability was significantly positive, we compared the within-stimulus values with the across-stimulus values for each frequency band (paired t-test with Bonferroni correction for 43 frequencies, α = 0.0012). In awake subjects we identified a restricted range of frequencies between 10.5 and 20 Hz where visual stimuli enhanced the phase reliability (Fig. 6g,i). In anesthetised animals, average PDI values were larger than in awake animals and all frequencies tested had single stream PDI values that were significantly non-zero (Fig. 6h,j). We therefore conclude that visual stimulation elicited reliable changes in the LFP phase in auditory cortex. In contrast to LFP phase, a parallel analysis of across trial power reliability showed no significant effect of visual stimuli on LFP power in any frequency band (Fig.S7a,c).

If visual information was only conveyed in the case of temporally coherent stimuli, this might indicate that the locus of binding was outside of auditory cortex and that the information being provided to auditory cortex already reflected an integrated auditory-visual signal. The LFP is thought to reflect the combined synaptic inputs to a region (Viswanathan and Freeman 2007) and so significant PDI values for both temporally independent and coherent stimuli suggests that the correlates of binding observed in auditory cortex were not simply inherited from its inputs. Since there were significant PDI values for both temporally independent and coherent stimuli, we next asked whether there were any frequencies at which phase coherence was significantly greater in AV stimuli which were temporally coherent compared to temporally independent. We performed a pairwise comparison of single stream PDI values obtained from temporally coherent and independent stimuli, for all frequency points. In awake animals, PDI values were similar for temporally coherent and temporally independent stimuli, except in the 10.5-12.5 Hz band where coherent stimuli elicited significantly greater phase coherence (Fig. 6k). In anaesthetised animals, the single stream PDI did not differ between coherent and independent stimuli at any frequency (Fig. 6l). Together these data suggests that visual inputs modulate the phase of the field potential in auditory cortex largely independently of any temporal coherence between auditory and visual stimuli. This finding supports the conjecture that multisensory binding occurs within auditory cortex.

To understand whether the same mechanisms could underlie the visual-stimulus induced enhancement of a temporally coherent sound in a mixture, we performed similar analyses on the data collected in response to the dual stream stimuli. We generated within-stimulus ITPC values for each dual-stream stimulus (i.e. A12V1 and A12V2) and across-stimulus ITPC by randomly selecting responses across visual conditions. We then expressed the difference as the dual stream phase dissimilarity index (dual stream PDI, Fig. 6m,n). Since the auditory components were identical in each dual stream stimulus, the influence of the visual component on LFP phase could be isolated as non-zero dual stream PDI values (paired t-test, Bonferroni corrected, α = 0.0012). In awake animals, the dual stream PDI was significantly non-zero at 10.5-12.5 (Fig.6m) whereas in anesthetised animals, we found positive dual stream PDI values across all frequencies tested (Fig.6n). In anesthetised animals, where we could use the responses of units to noise and light flashes to categorise units as auditory, visual or auditory-visual, we confirmed significant PDI values in the LFP recorded on the same electrode as units in each of these subpopulations (Fig.S3l). In awake animals, we tested auditory visual stimuli presented at three different modulation rates (7, 12 and 17Hz) and confirmed that significant PDI values were obtained at very similar LFP frequencies across these modulation rates - consistent with these being evoked phase alignments rather than stimulus-entrained oscillations (Fig. S7i). Additional evidence for that hypothesis comes from the fact that, in the awake data, the frequencies at which the single and dual stream PDI values are significant are entirely non-overlapping with the modulation rate of the stimulus, which was band-limited to 7 Hz.

### Visual cortex mediates visual-stimulus induced LFP changes in auditory cortex

Visual inputs to auditory cortex potentially originate from many sources: in the ferret, multiple visual cortical fields are known to innervate auditory cortex (Bizley et al., 2007), but frontal and thalamic areas are additional candidates for sources of top-down and bottom-up multisensory innervation. To determine the origin of the visual effects that we observe in auditory cortex, we performed an additional experiment in which we cooled the gyral surface of the posterior suprasylvian sulcus where visual cortical fields SSY (Cantone et al., 2006) and Area 21 (Innocenti et al., 2002) are located (Fig. 7a). Neural tracer studies have demonstrated that these areas directly project to auditory cortex in the ferret (Bizley et al., 2007). We used a cooling loop cooled to 9-10°C to reversibly silence neural activity within <500 μm of the loop (see Fig. 7b, see also Wood et al., 2017). Using simple noise bursts and light flashes at each site that we cooled, we verified that cooling visual cortex did not alter the response to noise bursts in auditory cortex (repeated measures ANOVA on spike rates in response to a noise burst pre-cooling, during cooling, after cooling, F(2,164) = 0.42 p=0.88), but did reversibly attenuate the spiking response to light flashes in visual cortical sites >500 μm from the cooling loop (repeated measures ANOVA F_(2,92)_ = 6.83 p=0.001, post-hoc comparisons shows pre-cool and cool-post were significantly different, pre-post were not significantly different indicating the effects were reversible) and under the loop (F(2,210) = 30.2586; p = 2.8350e-12, pre-cool vs. cooled, cooled vs. post-cooled significantly different, pre-post not significantly different). We measured responses to the single stream stimuli in auditory and visual cortex before and during cooling. From the LFP, we calculated the across-trial-phase coherence and phase dissimilarity indexes (as in Fig. 6). Cooling visual cortex significantly decreased the magnitude of the single stream PDI values in auditory cortex (Fig 7e,h). A 3-way repeated measures ANOVA with factors of visual condition (coherent/independent), frequency, and cortical temperature (warm/cooled) on the SS PDI values obtained in auditory cortex showed a main effect of frequency (F_(88,22605)_ = 47.91, p < 0.1*10^9^) and temperature (F_(1,22605)_ = 1072, p < 0.1*10^9^,) but not visual condition (p=0.49). In contrast, LFP at recording sites in visual cortex away (>500 μm) from the loop were unaffected by cooling (3-way ANOVA demonstrated that the magnitude of the SS PDI value was influenced by frequency (F_(88,17265)_ = 24.73, p<1*10^9^), but not temperature (p=0.75) or visual condition (p=0.29), Fig 7i-n). From these data, we conclude that the influence of visual stimuli on the auditory cortical field potential phase is mediated, at least in part, by inputs from visual cortical areas SSY and 21. While cooling does not allow us to confirm that visual inputs are direct mono-synaptic connections (Bizley et al., 2016a), the observation that the phase effects in other areas of visual cortex are unaffected suggests that cooling selectively influenced communication between auditory and visual cortices rather than supressing visual processing generally.

**Figure 7:**
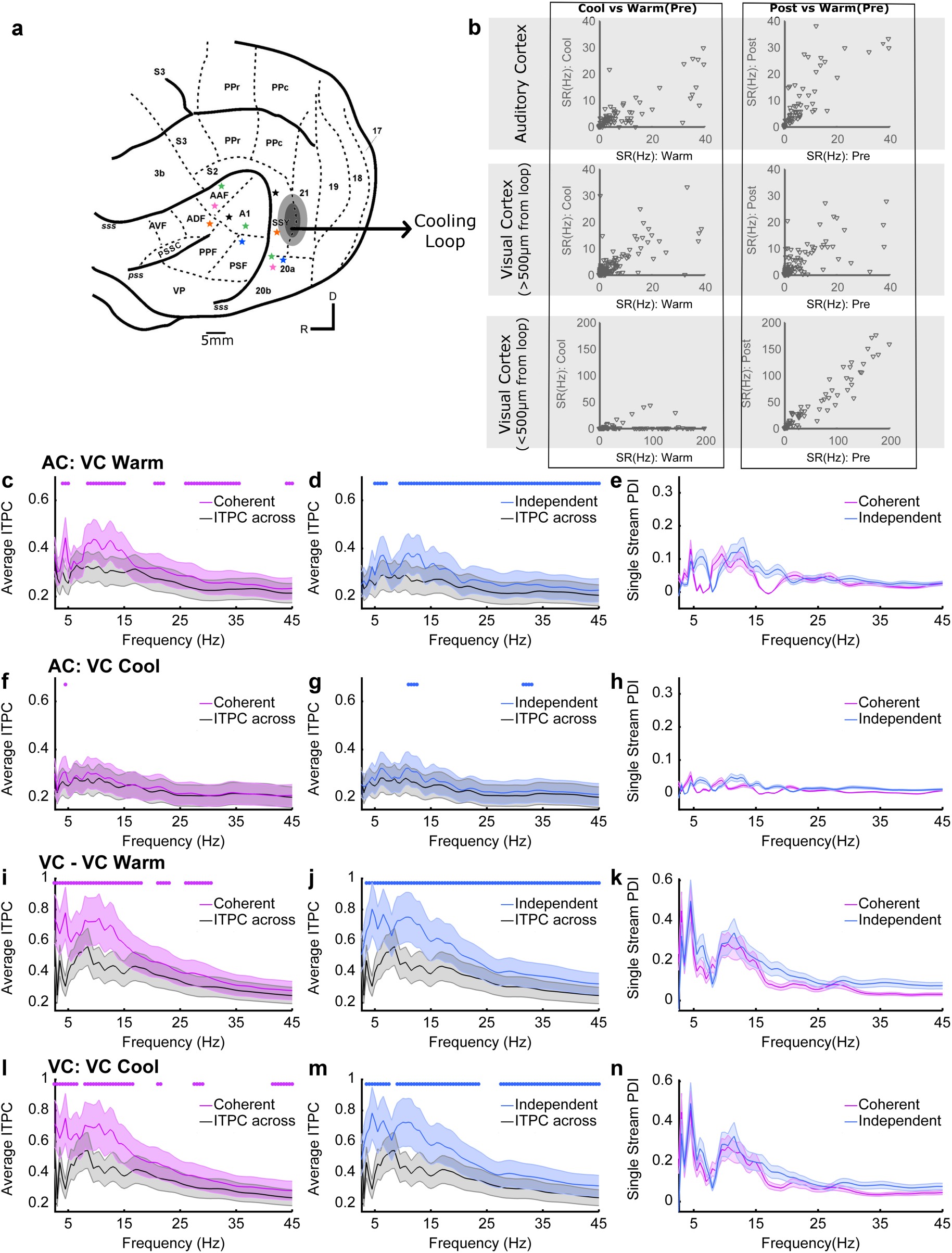
Visual stimulus induced LFP phase changes in auditory cortex are mediated in visual cortex. **a** Schematic showing the location of auditory cortical recording sites and the location of a cooling loop (black, grey line marks the 500 μm radius over which cooling is effective (Wood et al., 2017) which was used to inactivate visual cortex. Individual recording sites contributing to c-n are shown with stars (simultaneous recordings are marked in the same colour). **b** Spike rate responses in Auditory Cortex (top row) and visual cortex (bottom row, sites >500 μm from the loop) in response to noise bursts or light flashes before, during and after cooling. **c,d**, inter-trial phase coherence values for the coherent (**c**) and independent stimuli (**d**) AV stimuli recorded in auditory cortex (AC) prior to cooling visual cortex (VC) compared to the shuffled null distribution (ITPC across). Asterisks indicate the frequencies at which the ITPC values are significantly different from the shuffled ITPC-across distribution **e** single stream Phase Dissimilarity Index values calculated from the ITPC values in **c** and **d**. **f-h** - as c-e but while visual cortex was cooled to 9 degrees. **i-n** as c-h but for sites in visual cortex >500um from the cooling loop. c-h includes data from 83 sites from 6 electrode penetrations, i-n includes data from 47 sites from 5 penetrations.

## Discussion

Here we provide insight into how and where auditory-visual binding occurs, and provide evidence that this effect is mediated by cortico-cortical interactions between visual and auditory cortex. While numerous studies have reported the incidence of auditory-visual interactions in auditory cortex over the past decade (Bizley et al., 2007a; Chandrasekaran et al., 2013; Ghazanfar et al., 2005; Kayser et al., 2008; Kayser et al., 2010, Perrodin et al., 2015), evidence for the functional role has remained less apparent. Here we show that one role for the early integration of auditory and visual information is to support auditory scene analysis. Visual stimuli elicit reliable changes in the phase of the local field potential in auditory cortex, irrespective of auditory-visual temporal coherence, indicating that the inputs to auditory cortex reflect the unisensory stimulus properties. When the visual and auditory stimuli are temporally aligned, phase resets elicited by the visual stimulus interacts with feed-forward sound-evoked activity and results in spiking output that more precisely represents the temporally coherent sound within auditory cortex. These results are consistent with the binding of cross-modal information to form a multisensory object because they result in a modification of the representation of the sound which is not restricted to the features that link auditory and visual signals but extends to other non-binding features. These data provide a physiological underpinning for the pattern of performance observed in human listeners performing an auditory selective attention task, in which the detection of a perturbation in a stimulus stream is enhanced or impaired when a visual stimulus is temporally coherent with the target or masker auditory stream respectively (Maddox et al., 2015). The effects of the visual stimulus on the representation of an auditory scene can be observed in anesthetised animals suggesting that these effects can occur independently of attentional modulation.

Previous investigations of the impact of visual stimuli on auditory scene analysis have frequently used speech stimuli. Being able to see a talker’s mouth provides listeners with information about the rhythm and amplitude of the speech waveform which may help listeners by cueing them to pay attention to the auditory envelope (Peelle and Sommers, 2015) as well as by providing cues to the place of articulation that can disambiguate different consonants (Sumby and Pollack, 1954). However, the use of speech stimuli makes it difficult to dissociate general multisensory mechanisms from speech-specific ones when testing in human subjects. Therefore, in order to probe more general principles across both human (Maddox et al., 2015) and non-human animals (here), we chose to employ continuous naturalistic non-speech stimuli that utilized modulation rates that fell within the range of syllable rates in human speech but lacked any linguistic content. Previous work has demonstrated that a visual stimulus can enhance the neural representation of the speech amplitude envelope both in quiet and in noise (Crosse et al., 2015; Crosse et al., 2016; Luo et al., 2010,Park et al., 2016), but functional imaging methods make it difficult to demonstrate enhanced neural encoding of features beyond the amplitude envelope. The implication of our findings is that representation of the spectro-temporal features that allow speech recognition such as voice pitch would be enhanced in auditory cortex when a listener views a talker’s face, even though such spectro-temporal features may not be represented by the visual stimulus.

Visual speech information is hypothesised to be relayed to auditory cortex through multiple routes in parallel to influence the processing of auditory speech: Our data support the idea that early integration of visual information occurs (Möttönen et al., 2004; Okada et al., 2013; Peelle and Sommers, 2015; Schroeder et al., 2008) and is likely to reflect a general phenomenon whereby visual stimuli can cause phase-entrainment in the local field potential. Within this framework, cross-modal binding potentially results from the temporal coincidence of evoked auditory responses and visual-stimulus elicited inputs that we observe as phasic changes of the LFP.

Consistent with previous studies, our analysis of local field potential activity revealed that visual information reliably modulated LFP phase in auditory cortex (Chandrasekaran et al., 2013; Ghazanfar et al., 2005; Kayser et al., 2008; Perrodin et al., 2015). This occurred independently of the modulation frequency of the stimulus suggesting that, rather than entraining oscillations at the stimulus modulation rate, relatively broadband phase resets are triggered by particular features within the stream (presumably points at which the luminance changed rapidly from low-high amplitude). LFP reflects the synaptic inputs to a region and LFP phase synchronization is thought to arise from fluctuating inputs to cortical networks (Lakatos et al., 2007; Mazzoni et al., 2008; Szymanski et al., 2011). Since neuronal excitability varies with LFP phase (Jacobs et al., 2007; Klimesch et al., 2007; Lakatos et al., 2013; Lőrincz et al., 2009), synaptic inputs from visual cortex may provide a physiological mechanism through which temporally coincident cross-sensory information is integrated. Our analysis allowed us to isolate changes in LFP phase that were directly attributable to the visual stimulus and identify reliable changes in LFP phase irrespective of whether the visual stimulus was temporally coherent with the auditory stimulus. Such results suggest that the observed effects of cross-modal temporal coherence were not simply inherited within the inputs to auditory cortex. Moreover the effects that we observed in the LFP were lost when we silenced visual cortex, indicating that inputs from visual cortex are a key contributor to the effects of auditory-visual temporal coherence that we observed in auditory cortex. Our finding that visual stimulation elicited reliable phase modulation in both awake and anesthetised animals suggests that bottom-up cross-modal integration interacts with selective attention, which has also been associated with modulation of phase information in auditory cortex (Golumbic et al., 2013, Park et al., 2016). While our data suggest that cross-modal binding can occur in the absence of attention, it is likely that the additional neural pathways engaged during selective attention act to further enhance the representation of attended cross-modal objects.

In both awake and anesthetised animals we observed that visual stimuli elicit robust effects on the LFP phase, that auditory-visual temporal coherence shapes the response to a sound mixture such that temporally coherent auditory-visual stimuli are more reliably represented in the spiking response, and that the spiking response to auditory timbre deviants (a non-binding feature) was enhanced. While these key findings were recapitulated in both states, there were some important differences: Firstly, in the awake animal the phase alignment in the LFP was generally smaller in magnitude and was only significantly modulated across a smaller range of frequencies (10.5-20 Hz as opposed to 4-45 Hz). Such differences are consistent with a dependence of oscillatory activity on behavioural state (Tukker et al., 2007; Voloh and Womelsdorf, 2016; Wang, 2010). Secondly, in the awake animal we observed a significant increase in the phase reliability (at 10.5-12.5 Hz) for temporally coherent auditory-visual stimuli when compared to temporally independent stimuli. Since the neural correlates of multisensory binding are evident in the anesthetised animal, the specific increase in alpha phase reliability that occurred only in awake animals in response to temporally coherent auditory-visual stimulus pairs (Fig. 6k,m) may indicate an attention-related signal triggered by temporal coherence between auditory and visual signals, or an additional top-down signal conveying cross-modal information. Phase resetting or synchronisation of alpha phase has been associated both with enhanced functional connectivity (Voloh and Womelsdorf, 2016) and as a top-down predictive signal for upcoming visual information (Samaha et al., 2015). Understanding how attention engages additional brain networks and disambiguating these possibilities would require simultaneous recordings in auditory and visual cortex recording while trained animals performed the auditory selective attention task which motivated this study. Finally, in awake and anesthetised animals we observed that the impact of auditory-visual temporal coherence on the representation of sound mixtures (as assessed by visual preference scores) was of a similar magnitude in the primary areas (A1 and AAF, located in the MEG). In contrast, in the awake animal, neurons in the PEG, where secondary tonotopic fields PPF and PSF are located, had significantly higher VPI scores than those in the MEG, while in anesthetised animals VPI scores were statistically indistinguishable across cortical fields. This suggests that in the awake animal additional mechanisms exist to enhance the effects that are present in the primary areas. These results were mirrored in the impact of auditory-visual temporal coherence on non-binding features (as assessed by the impact of auditory-visual temporal coherence on deviant detection ability) where the visual stimulus had a stronger influence in PEG than MEG in the awake animal, and did not differ across regions (and was overall of a smaller magnitude) in anesthetised animals. Our cooling studies (in anesthetised animals) do not allow us to determine whether this enhancement reflects the greater variety of inputs from visual cortex that terminate in secondary as opposed to primary auditory cortex (Bizley et al., 2007), top down inputs from higher areas (e.g. parietal or frontal cortex), or are a consequence of intracortical processing within auditory cortex.

Temporal coherence between sound elements has been proposed as a fundamental organising principle for auditory cortex (Elhilali et al., 2009; O'Sullivan et al., 2015b) and here we extend this principle to the formation of cross-modal constructs. Our data provide evidence that one role for the early integration of visual information into auditory cortex is to resolve competition between multiple sound sources within an auditory scene and that these neural computations occur pre-attentively. While some proponents of a temporal coherence based model for auditory streaming have stressed the importance of attention in auditory stream formation (Lu et al., 2017), neural signatures of temporal-coherence based streaming are present in passively listening subjects (O'Sullivan et al., 2015a; Teki et al., 2016). Previous studies have demonstrated a role for visual information in conveying lip movement information to auditory cortex (Chandrasekaran et al., 2013; Crosse et al., 2015; Ghazanfar et al., 2005; Golumbic et al., 2013), but such stimuli make it difficult to separate sensory information from linguistic cues. Our data obtained using non-speech stimuli provide evidence that at least part of the boost provided by visualising a speaker’s mouth arises from a more general (language-independent) phenomenon whereby visual temporal cues facilitate auditory scene analysis through the formation of cross-sensory objects. Our data are supportive of visual cortical areas providing at least one source of information. Other visual cortex fields and sub-cortical structures innervate tonotopic auditory cortex (Bizley et al., 2007; Budinger et al., 2006) and may potentially provide additional visual inputs to auditory cortex. Further dissecting the origin of visual innervation requires experiments that allow pathway specific manipulation of neuronal activity (for example by silencing the terminal fields of neurons that project from a candidate area into auditory cortex, Bizley et al., 2016a).

Finally, the neural correlates of multisensory binding were apparent in units which best discriminated either the auditory or visual characteristics of single auditory-visual streams, although the magnitude of the effects was stronger in visual-discriminating units. Nevertheless, both classes of neurons showed enhanced encoding of temporally coherent versus temporally independent auditory visual streams suggesting that both subgroups could be described as auditory-visual. This was confirmed in anesthetised animals where neurons were additionally characterised with simple stimuli (noise bursts and light flashes) and revealed that 54% of visual-discriminating stimuli and 41% of auditory-discriminating neurons were classified as auditory-visual. Together, these results suggest that multisensory processing is prevalent throughout auditory cortex and that cross-sensory processing has the potential to have a significant impact on the representation of acoustic features in auditory cortex.

In summary, activity in auditory cortex was reliably affected by visual stimulation in a manner that enhanced the representation of temporally coherent auditory information. Enhancement of auditory information was observed for sounds presented alone or in a mixture and for sound features that were related to (amplitude) and orthogonal to (timbre) variation in visual input. Such processes provide mechanistic support for a coherence-based model of cross-modal binding in object formation and indicate that one role for the early integration of visual information in auditory cortex is to support auditory scene analysis.

## Acknowledgments

This work was funded by grants to each author: JKB: Wellcome Trust / Royal Society WT098418MA; Biotechnology and Biological Sciences Research Council (BB/H016813/1), and an Action on Hearing Loss Studentship (596: UEI: JB); RKM: NIH R00DC014288 and Hearing Health Foundation Emerging Research Grant; AKCL: NIH R01DC013260; and an International Exchanges Scheme award from the Royal Society to JKB and AKCL.

## Author contributions

HA, RKM, AKCL, JKB Conception and design, HA, SMT, KCW, GPJ, JKB Acquisition of data, HA, JKB Analysis and interpretation of data, HA, SMT, RKM, AKCL, JKB Drafting or revising the article.

## Methods

### CONTACT FOR REAGENT AND RESOURCE SHARING

Further information and requests for resources and reagents (data and MATLAB code) should be directed to and will be fulfilled by the Lead Contact, Jennifer Bizley.

### EXPERIMENTAL MODEL DETAILS

The experiments were approved by the Animal Welfare and Ethical Review Board of University College London and The Royal Veterinary College, and performed under license from the UK Home Office (PPL 70/7267) and in accordance with the Animals Scientific Procedures Act 1986.

Neural responses were recorded in a total of 19 awake pigmented adult female ferrets (Mustela putorius furo; 1-5 years old). Fourteen of these animals contributed data to the awake dataset: Data from 9 of these animals was used for the main experiment (532 units), data from 11 other animals (6/9 in the main experiment, plus five other ferrets, totalling 128 units) was collected for additional control analysis (Fig. 3e, Fig. S5). Females (700-1500g, wild type) were co-housed in groups of 2-9. These animals were trained in a variety of psychoacoustic tasks unrelated to the current study prior to and after the implantation of recording electrodes. Animals were tested for this study on days when they were not participating in psychoacoustic testing. Five adult females were used to record responses under anaesthesia.

## METHOD DETAILS

### Animal preparation

Full methods for recording under anaesthesia can be found in Bizley et al., (2009). Briefly, ferrets were anesthetized with medetomidine (Domitor; 0.022mg/kg/h; Pfizer, Sandwich, UK) and ketamine (Ketaset; 5mg/kg/h; Fort Dodge Animal Health, Southampton, UK). The animal was intubated and the left radial vein was cannulated in order to provide a continuous infusion (5 ml/h) of a mixture of medetomidine and ketamine in lactated ringers solution augmented with 5% glucose, atropine sulfate (0.06 mg/kg/h; C-Vet Veterinary Products) and dexamethasone (0.5 mg/kg/h, Dexadreson; Intervet, UK). The ferret was placed in a stereotaxic frame in order to implant a bar on the skull, enabling the subsequent removal of the stereotaxic frame. The left temporal muscle was largely removed, and the suprasylvian and pseudosylvian sulci were exposed by a craniotomy, revealing auditory cortex (Kelly et al., 1986). The dura was removed over auditory cortex and the brain protected with 3% agar solution. The eyes were protected with zero-refractive power contact lenses. The animal was then transferred to a small table in a sound-attenuating chamber. Body temperature, end-tidal CO2, and the electrocardiogram were monitored throughout the experiment. Experiments typically lasted between 36 and 56 h. Neural activity was recorded with multisite silicon electrodes (Neuronexus Technologies) in a 1x 16, 2x 16 or 4x 8 (shank x number of sites) configuration. For experiments in which visual cortex was cooled we extended the craniotomy caudally to expose visual cortex and placed a cooling loop over the posterior suprasylvian gyrus. Details of the manufacture of the cooling loop and validation of its efficacy in the ferret animal model are provided in full in (Wood et al., 2017).

Full surgical methods for recording implanting electrode arrays to facilitate recording from awake animals are available in Bizley et al. (2013). Briefly, animals were bilaterally implanted with WARP-16 drives (Neuralynx, Montana, USA) loaded with high impedance tungsten electrodes (FHC, Bowdoin, USA) under general anaesthesia (medetomidine and ketamine induction, as above, isoflurane maintenance 1-3%). Craniotomies were made over left and right auditory cortex, a small number of screws were inserted into the skull for anchoring and grounding the arrays, and the WARP-16 drive was anchored with dental acrylic and protected with a capped well. Recording electrodes in awake animals targeted tonotopic auditory cortex (area MEG, containing fields A1 and AAF, and PEG, tonotopic belt areas PPF and PSF are located). Auditory fields were estimated prior to implantation based on known sulcal landmarks and confirmed with regular assessments of frequency tuning and post-mortem histology. Animals were allowed to recover for a week before the electrodes were advanced into auditory cortex. Pre-operative, peri-operative and post-operative analgesia were provided to animals under veterinary advice. Recordings were made over the next 1-2.5 years, with electrodes individually advanced every few weeks until the thickness of auditory cortex was traversed. Recordings were made while animals were passively listening/watching stimuli and holding their head at a water spout. During the recording a continuous stream of water was delivered from the spout.

#### Stimulus Presentation

All stimuli were created using TDT System 3 hardware (Tucker-Davis Technologies, Alachua, FL) and controlled via MATLAB (Mathworks, USA). For recordings in awake animals, sounds were presented over two loud speakers (Visaton FRS 8). Water deprived ferrets were placed in a dimly lit testing box (69 × 42 × 52 cm length × width × height) and received water from a central reward spout located between the two speakers. Sound levels were calibrated using a Brüel and Kjær (Norcross, GA) sound level meter and free-field ½-inch microphone (4191). Auditory streams were presented at 65 dB SPL (Fig. 1a). Visual stimuli were delivered by illuminating the spout with a white LED which provided full field illumination (Precision Gold N76CC Luxmeter, 0 to 36.9 lux). The animals were not required to do anything other than maintain their heads in position at the spout where they were freely rewarded. Recording was terminated when animals were sated.

For anesthetised recordings, acoustic stimuli were presented using Panasonic headphones (Panasonic RP-HV297, Bracknell, UK) at 65 dB SPL. Visual stimuli were presented with a white Light Emitting Diode (LED) which was placed in a diffuser at a distance of roughly 10 cm from the contralateral eye so that it illuminated virtually the whole contralateral visual field.

#### Stimuli and data acquisition

*Auditory stimuli* were artificial vowel sounds that were created in Matlab (MathWorks, USA). In the behavioural experiment that motivated this study(Maddox et al., 2015), stimuli were 14 seconds in duration. However, we adapted the stimulus duration in awake recordings to 3 seconds in order to collect sufficient repetitions of all stimuli, and to ensure animals maintained their head position facing forwards for the whole trial duration. The animals were observed constantly via a webcam and recording was terminated / paused if the animal’s head moved from the centre spout. In the anesthetised recording stimulus streams were 14 seconds long, as in the human psychophysics but we only analysed the first 3 seconds to ensure datasets were directly comparable (see also Fig. S7 e-h which replicates analysis for 3 second and 14 second stimuli).

Stimulus A1 was the vowel [u] (formant frequencies F1-4: 460, 1105, 2857, 4205 Hz, F0= 195Hz), A2 was [a] (F1-4: 936, 1551, 2975, 4263 Hz, F0= 175Hz). Streams were amplitude modulated with a noisy lowpass (7 Hz cut-off) envelope. Unless specifically noted, the timbre of the auditory stream remained fixed throughout the trial. However, we also recorded responses to auditory streams that included brief timbre deviants. As in our previous behavioural study, deviants were 200ms epochs in which the identity of the vowel was varied by smoothly changing the first and second formant frequencies to and from those identifying another vowel. Stream A1 was morphed to/from [ε] (730, 2058, 2857, 4205 Hz) and A2 to/from [i] (437, 2761, 2975, 4263 Hz).

*Visual stimuli* were generated using an LED whose luminance was modulated with dynamics that matched the amplitude modulation applied to A1 or A2. In single stream conditions a single auditory and single visual stream were presented (e.g. A1V1, A1V2, A2V1, or A2V2) whereas in dual stream conditions both auditory streams were presented simultaneously, accompanied by a single visual stimulus (A12V1, A12V2, A12V1 A12V2) (Fig. 1e). Auditory streams were always presented from both speakers so that spatial cues could not facilitate segregation, and stimulus order was varied pseudo-randomly. In the anesthetised recordings each stimulus was presented 20 times. In the awake dataset, where recording duration was determined by how long the ferret remained at the central location (mean repetitions: 20, minimum: 14, maximum: 34).

During anaesthetised recordings, pure tone stimuli (150 Hz to 19 kHz in 1/3-octave steps, from 10 to 80 dB SPL in 10 dB, 100 ms in duration, 5 ms cosine ramped) were also presented. These allowed us to characterize individual units and determine tonotopic gradients, so as to confirm the cortical field in which any given recording was made. Additionally broadband noise bursts and diffuse light flashes (100 ms duration, 70 dB SPL) were presented and used to classify a stimulus as auditory, visual or auditory visual. LFPs were subjected to current source density analysis to identify sources and sinks as described by Kaur et al. (2005).

### Cortical cooling

During these experiments we made joint recordings in visual cortex (usually >500 μm from the cooling loop, in order to determine whether visual cortical processing was impaired generally) and auditory cortex simultaneously. We recorded responses to the single stream stimuli before and during cooling, and, at each site additionally recorded responses to noise bursts and light flashes before, during and after cooling. We used the responses to simple stimuli such as these to show that we could recover the original spiking responses (and data was excluded from any recording sites in which did not return to within 20% (a common criterion used in cooling studies: e.g. Antunes and Malmeirca, 2011) of their pre-cooling spike rates). We did not record responses to the longer stimuli used in this study in the post-cooling condition as the additional recording time for these stimuli would have compromised our ability to record across several different sites in each animal.

## QUANTIFICATION AND STATISTICAL ANALYSIS

Electrophysiological data were analysed offline. Spiking activity and local field potential signals were extracted from the broadband voltage waveform by filtering at 0.3-5kHz and 1-150 Hz respectively. Spikes were detected, extracted and then sorted with a spike-sorting algorithm (WaveClus, Quiroga et al., 2004).

### Spiking responses

We used a Euclidean distance based pattern classifier (Foffani and Moxon, 2004) with leave-one-out cross validation to determine whether the neuronal responses to different stimuli could be discriminated. Spiking responses to a given stimulus were binned into a series of spike counts from stimulus onset (0 s) to offset (3s) in 20 ms bins. The average across-repetition response to each stimulus (excluding the to-be-classified response) were used as templates and the response to a single stimulus presentation was classified by calculating the Euclidean distance between itself and the template sweeps and assigning it to the closest template. To determine whether the classifier performed significantly better than expected by chance, a 1000 iteration permutation test was performed where trials were drawn (with replacement) from the observed data and randomly assigned to a stimulus that was then used for template formation / decoding. A neural response was considered to be significantly informative about stimulus identity if the observed value exceeded the 95th percentile of the randomly-drawn distribution.

This approach allowed us to classify units according to their functional properties: auditory units discriminated two auditory stimuli based on the amplitude modulation of sound (A1 versus A2) regardless of visual dynamics, (Fig. 5a, b), visual units discriminated visual presentations based on temporal envelope of visual stimuli (V1 versus V2) regardless of auditory presentation (Fig. 5c,d) and AV units could do both. This approach was extended to classify dual stream responses by using the average response to each of the temporally coherent single stream stimuli (A1V1 or A2V2) as templates (Fig. 2,3, S2-4). Performance was (arbitrarily) expressed as the proportion of responses classified as being from the A1, and compared for the two dual stream stimuli with different visual conditions (Figure 5). All units in which either auditory or visual stimulus identity could be decoded were included in the dual-stream analysis. A VPI was derived from this measure as the difference between the percentage of A12V2 trials labelled A1 and the percentage of A12V1 trials labelled A1 (multiplied by -1 to make the index positive for units that were strongly influenced by the visual stimulus). Therefore units which were fully influenced by the identity of the visual stimulus would have a visual preference score of 100, while those in which the visual stimulus did not influence the response at all would have a score around 0 (Fig. 2e,h). We then assessed the significance of observed VPI scores using a permutation test (p < 0.05) in which the identity of single stream trials used to generate classifier templates was shuffled and the VPI recalculated for 1000 iterations.

#### Timbre deviant analysis

In order to determine how a visual stimulus influenced the ability to decode timbre deviants embedded within the auditory streams we used the cross-validated pattern classifier described above for analysing single stream stimuli to discriminate deviant from no-deviant trials. Responses were considered over the 200 ms time window that the deviant occurred (or the equivalent time point in the no-deviant stimulus) binned with a 10 ms resolution. Significance was assessed by a 1000 iteration permutation test in which trials were randomly drawn with replacement from deviant and no-deviant responses. The discrimination score was calculated as the proportion of correctly classified trials.

### Classification as auditory or visual with simple stimuli

During recordings made under anaesthesia, we also recorded responses to noise bursts and light flashes (both 100 ms duration) presented separately and together to compare how the proportion of auditory / visual discriminating units measured to naturalistic dynamic stimuli compared to more traditional artificial stimuli. Specifically, responsiveness was defined using a two-way ANOVA (factors: auditory stimulus [on/off] and visual stimulus [on/off]) on spike counts measured during stimulus presentation. We defined units as being sound-driven (main effect of auditory stimulus, no effect of visual stimulus or interaction), light-driven (main effect of visual stimulus, no effect of auditory stimulus or interaction) or auditory-visual (main effect of both auditory and visual stimuli or significant interaction; p < 0.05) as in (Bizley et al., 2007).

#### Phase/power dissimilarity analysis

Local field potential recordings were considered for all sites at which there was a significant driven spiking response, irrespective of whether that response could discriminate auditory or visual stream identity. For the single stream trials, we computed a single stream Phase Dissimilarity Index (PDI), which characterizes the consistency and uniqueness of the temporal phase/power pattern of neural responses to continuous auditory stimuli (Luo and Poeppel, 2007a). This analysis compares the phase (or power) consistency across repetitions of the same stimulus with a baseline of phase-consistency across trials in which different stimuli were presented.

In the first stage of PDI analysis, we obtained a time-frequency representation of each response using wavelet decomposition with complex 7-cycle Morlet wavelets in 0.5 steps between 2.5–45 Hz, resulting in 86 frequency points. Next, we calculated the inter-trial phase-coherence value (ITPC; Equ.1) at each time-frequency point, across all trials in which the same stimulus was presented. For each frequency band, the ITPC time-course was averaged over the duration of the analysis window and across all repetitions to obtain the average *within-stimulus ITPC*.

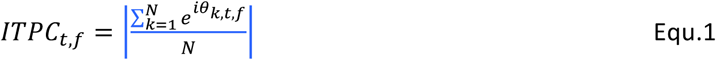

In which N is equal to the number of trials, and *θ* is the phase of trial *k* at a given frequency (*f*) and time (*t*). The *across-stimuli ITPC* was estimated using the same approach but using shuffled data, such that the ITPC was computed across trials with the same auditory stimulus but randomly drawn visual stimuli. The single stream phase dissimilarity index (Single stream PDI) was computed as the difference between the ITPC value calculated for *within* trials and the ITPC values calculated *across* visual trials (Equ.2). y. The dissimilarity function for each frequency bin *i* was defined as;

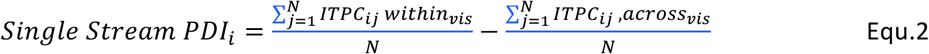

Large positive PDI indicate that responses to individual stimuli have a highly consistent response on single trials. Single stream PDI values were calculated for each stimulus type and then averaged across stimuli to calculate values for temporally coherent and temporally independent auditory visual stimuli. Single stream PDI was positive if within stimulus ITPC was larger than across-stimulus ITPC (pairwise t-test, p < 0.05 Bonferroni correction for 86 frequencies points) and was considered significant if a minimum of 2 adjacent bins exceeded the corrected threshold. PDI magnitude values were calculated by summing the PDI values across all significant frequencies.

Dual stream phase dissimilarity index (dual stream PDI) values were calculated by extending this approach for dual stream stimuli with the goal of determining how the temporal envelope of the visual stimulus influences the neural response to a sound mixture. To this end, we calculated the *within-dual ITPC* from the A12V1 trials and A12V2 trials separately and *across-dual ITPC* by randomly selecting trials from both stimuli (Equ.3). The within-dual and across-dual ITPCs were then averaged over time and subtracted to yield the dual stream PDI (Equ.3).

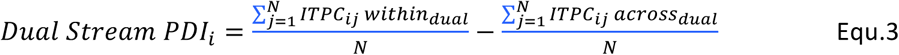

Positive dual stream PDI values indicate that the time course of the neural responses was influenced by visual input, despite the identical acoustic input. We determined whether the dual stream PDI was greater if the *within_dual ITPC* was significantly larger than *across_dual ITPC* (pairwise t-test, p < 0.05 Bonferroni correction, as above). PDI magnitude values were calculated by summing the PDI values across all significant frequencies.

### Analysis of responses during cooling

Our physiological recordings confirm that within the vicinity of the loop the inactivation spans all cortical layers. As the temperature change drops off with distance, at distances further from the loop the cooling is more restricted to superficial layers. These data are presented in full in Wood et al,. 2017.

## Supplemental Results

**Supplemental Figure 1 (related to Figure 1).**
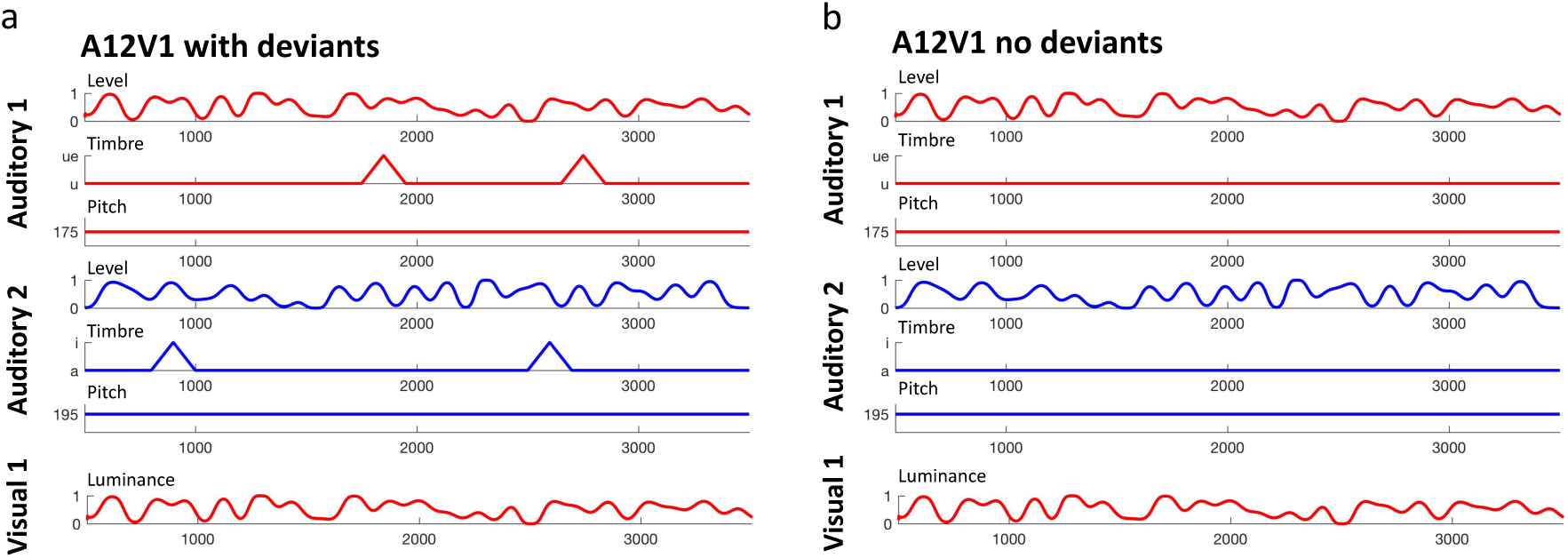
Schematic of the stimuli used in this study to illustrate the difference between stimuli with deviants (a) and without (b). Each row depicts the time course of each feature within the stimulus over a single trial. Importantly, the timing of the timbre deviants is not predicted by the temporally coherent changes in the binding features: here sound level and visual luminance.

**Supplemental Figure 2 (related to Figure 2).**
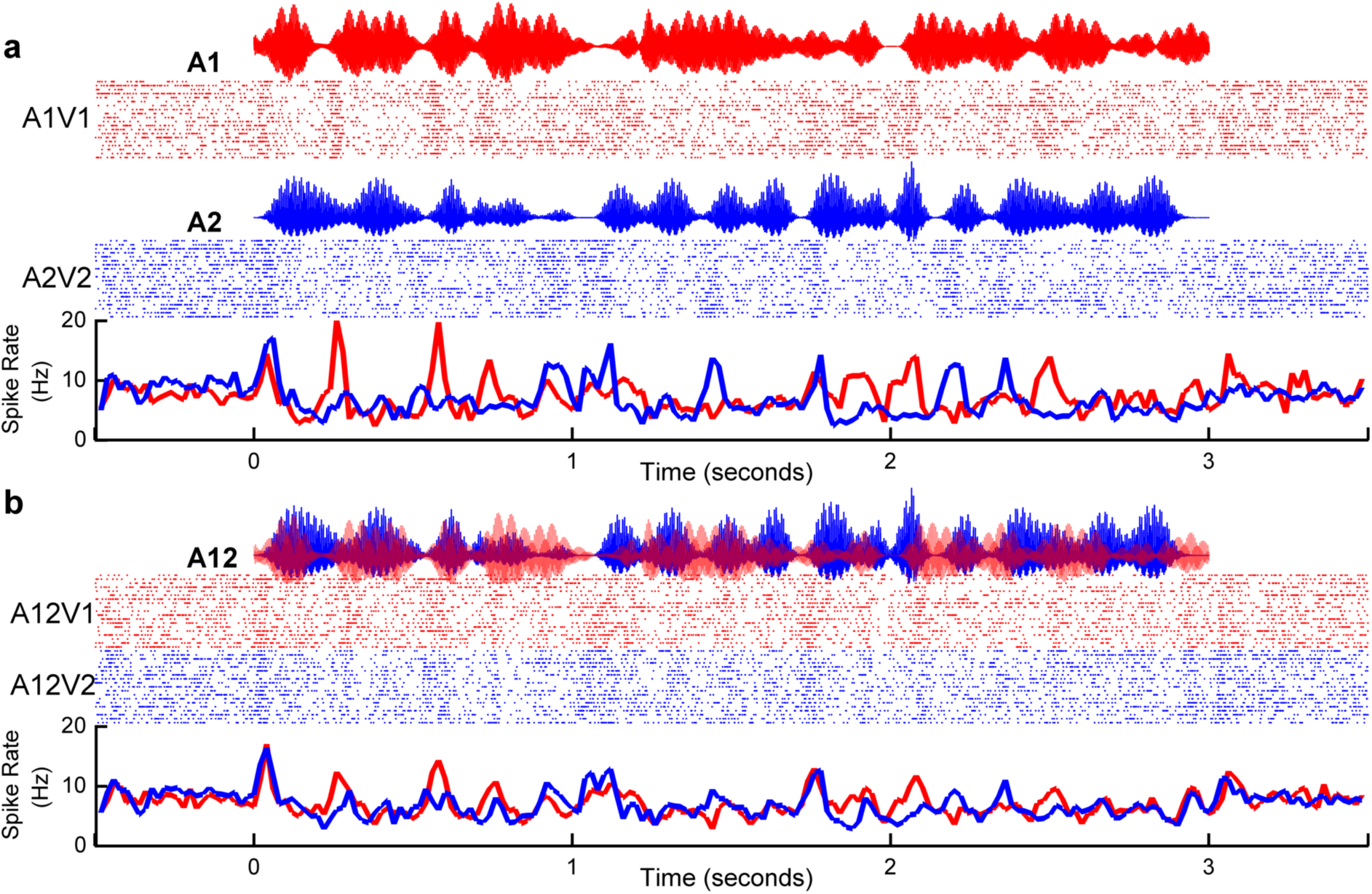
Visual stimuli can determine which sound stream auditory cortical neurons follow in a mixture: example auditory-discriminating unit. The spiking responses of an example unit are shown to coherent single stream (**a**) and dual stream stimuli (**b**). This example unit was an auditory discriminating unit recorded in an awake animal. In this example 68% (15/22) of responses were classified as A1 when the visual stimulus was V1, and 40 % of responses (9/22) were classified as A1 when the visual stimulus was V2, yielding a VPI score of 28%.

**Supplemental Figure 3 (related to Figures 2 and 3).**
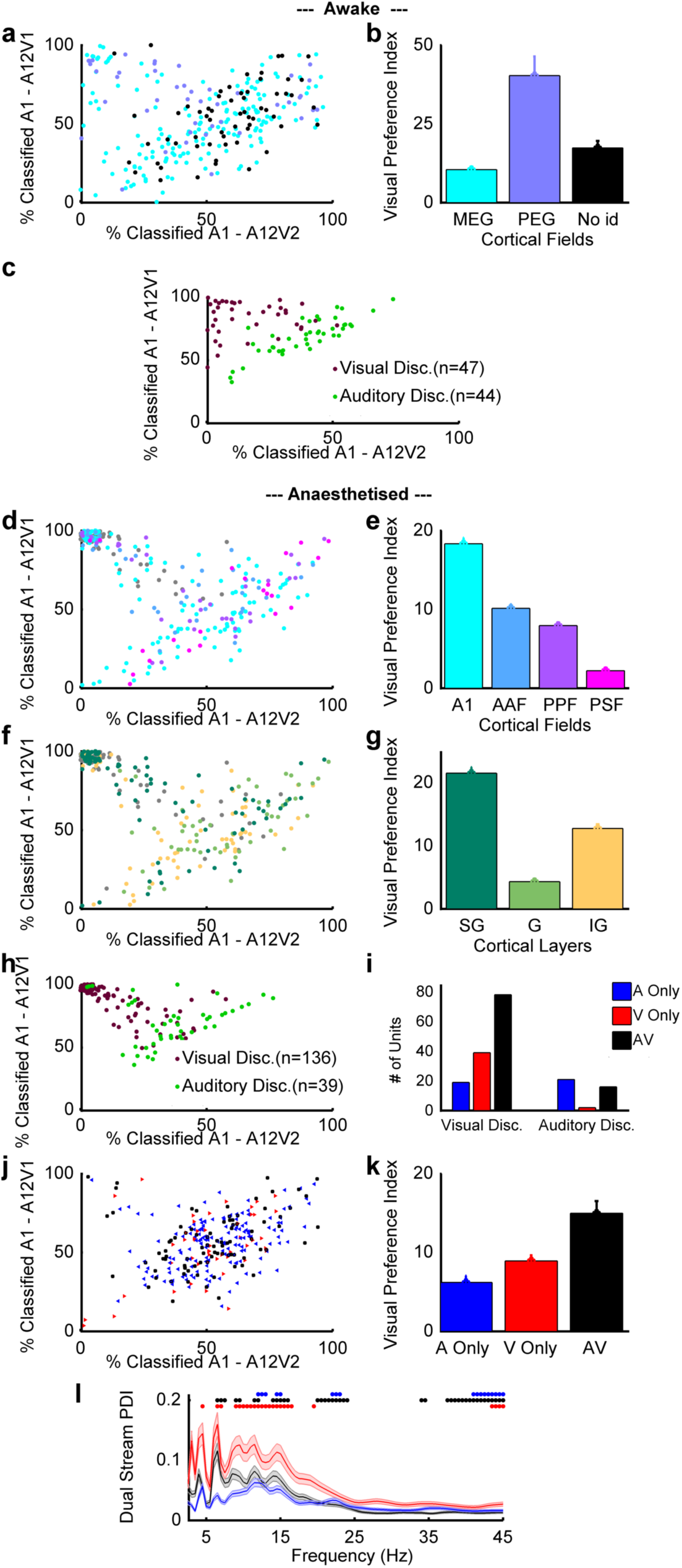
Effects of cortical field, cortical lamina and response type on visual modulation of dual stream responses. **a** The distribution of decoding values for dual stream responses, according to the visual condition (as in Figure 2C) colour coded according to recording location. Awake recordings were made from animals in which recording electrodes targeted the MEG (where A1 and AAF are located) and PEG (where fields PSF and PPF are located). The sampling density of our recording arrays (16 electrodes in a 4 × 4 grid with electrodes separated by 800 μm) does not provide the high spatial resolution necessary to determine recording locations precisely – particularly at the low frequency reversal that separates PEG from MEG in the ferret. Therefore some recording electrodes are classified as ‘unidentified’ as in these cases neither the frequency tuning, nor the post-mortem histology, allowed us to unambiguously ascribe the recording site to MEG or PEG. **b** VPI scores from **a** by cortical area (mean ±SEM). A one way ANOVA revealed there to be a significant effect of field (F(2,278) = 7.1354, p = 9.5072e-04), with post-hoc comparisons indicating PEG was significantly higher than MEG or unidentified sites. **c** re-plots the data in **a** showing only units with a significant VPI, colour-coded according to whether they were visual classified or auditory classified. **d,e,** as **a,b** but for the anesthetised dataset where we were able to make sufficient penetrations (30-50) to generate a high resolution tonotopic map and hence ascribe recording sites to cortical subfields. A one way ANOVA revealed there to be no significant effect of field on VPI (F(3,1130) = 2.1886, p =0.0877). Recordings in anesthetised animals were made with linear shank electrodes, facilitating current source density analysis to identify the cortical layers. **f** shows the distribution of decoding values in the dual stream condition according to recording location and depth in the anesthetised dataset. **g** summarises the data in **c** by cortical field. A one-way ANOVA across cortical layers showed a significant effect of layer (F(2,1134) = 3.1543, p= 0.0430) with post-hoc comparisons indicating that the VPI scores were greater in the supra-granular than granular layers. **h** plots the distribution of dual stream decoding values for only units with a significant VPI, colour-coded according to whether they were classified as auditory-discriminating or visual-discriminating. In the anesthetised animal we additionally used simple noise bursts and light flashes to describe units as auditory (A; n=160 units), visual (V, n=53) or auditory visual (AV; grey, n=94). **i** shows the distribution of A, V and AV units that were also classified as auditory-discriminating or visual-discriminating. Of 136 visual discriminating units with a significant VPI, 19 were categorised as auditory, 39 as visual and 78 as auditory-visual, of 39 auditory-discriminating units with significant VPI values 21 were auditory, 2 were visual and 16 were auditory visual Fig. S3i. **j,l,** as **d**,**e,** but with units colour coded according to whether they were classified as A, V or AV with simple stimuli. **k** mean (± SEM) dual stream phase dissimilarity index (PDI) values for recording sites categorised according to the spiking responses recorded there. Symbols indicate the frequencies at which the dual stream PDI index was significant (pairwise t-test, p < 0.001 with correction). While the phase effects are greatest at the sites where visual activity was recorded, significant dual stream PDI values were observed in all three unit types. In all three cases significant phase coherence was seen at 12Hz, 13.5Hz-14.5Hz and 42.5-44.5Hz. Modulation at 10-12 Hz was only observed at sites in which AV and V responses were recorded.

**Supplementary figure 4: Related to Figures 2,3 and 5.**
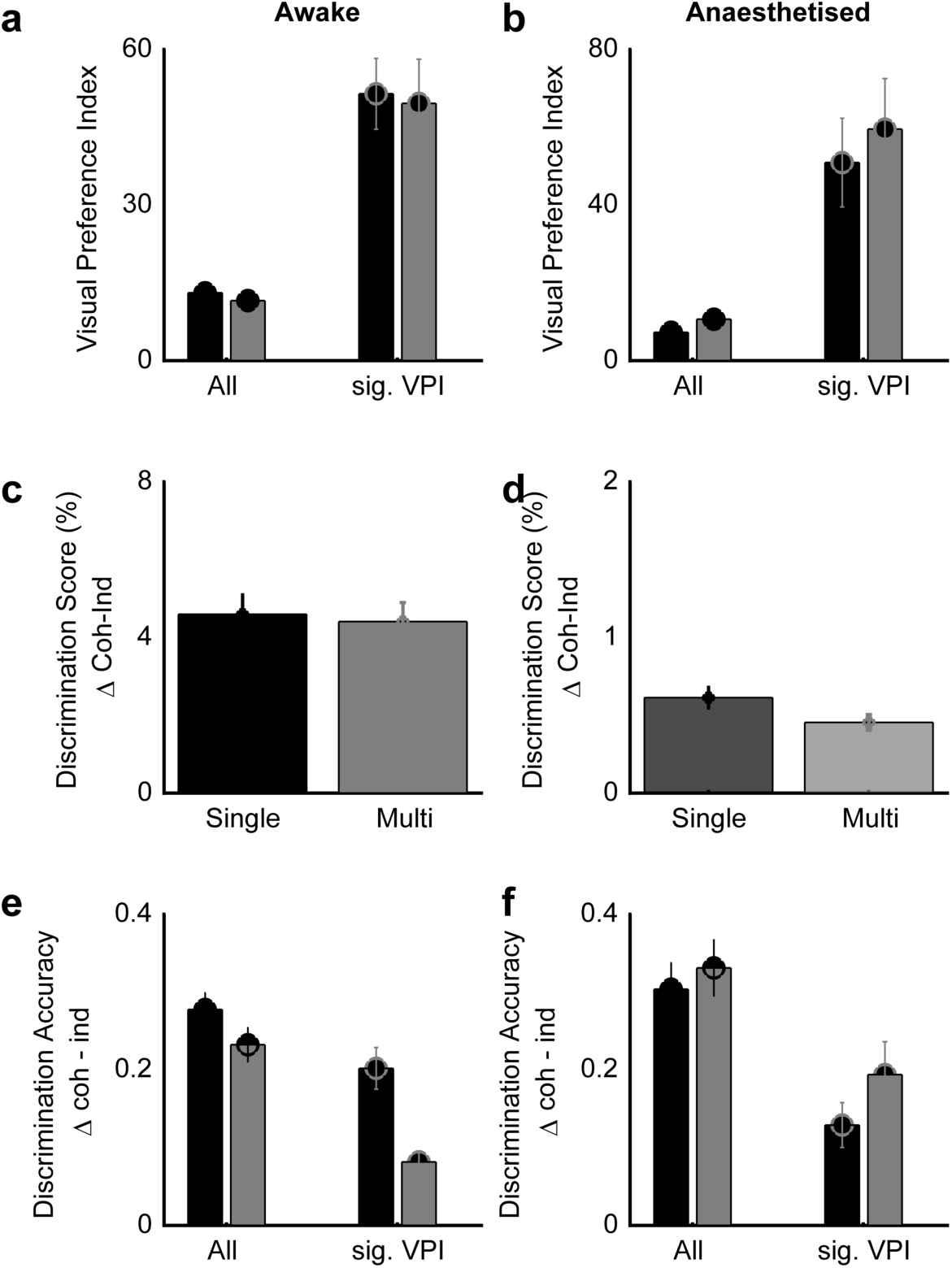
The effects of temporal coherence on single stream decoding and of visual identity on dual stream decoding are evident in both single and multi-units. **a,b** No effect of unit type (single versus multi-unit) was found for VPI values in awake recordings (all units: F(1,270) = 0.1595, p = 0.6899; significant VPI: F(1,90) = 0.1048, p = 0.7469) and anaesthetised recording (all units: F(1,332) = 0.4740, p = 0.4921; significant VPI: F(1,174) = 0.6867, p = 0.4123) **c,d** Discrimination scores for timbre deviant detection in dual stream stimuli is indistinguishable for single units and multi units in awake recording F(1,167) = 0.0326, p = 0.8570) and in anaesthetised recording (F(1,221)=0.8339, p = 0.3625) **e,f** Single units had significantly higher influence of temporal coherence on discrimination accuracy for single stream stimuli (as in Figure 5e,f) in awake recordings (All units: F(1,270) = 4.9916, p = 0.0263; for units with significant VPI: F(1,90) = 10.1780, p = 0.0020). Single and multiunits had equivalent performance in the anaesthetised dataset (F (1,332) = 1.2558, p = 0.2641; significant VPI: F(1,174) = 1.5121, p = 0.2262).

**Supplemental Figure 5.**
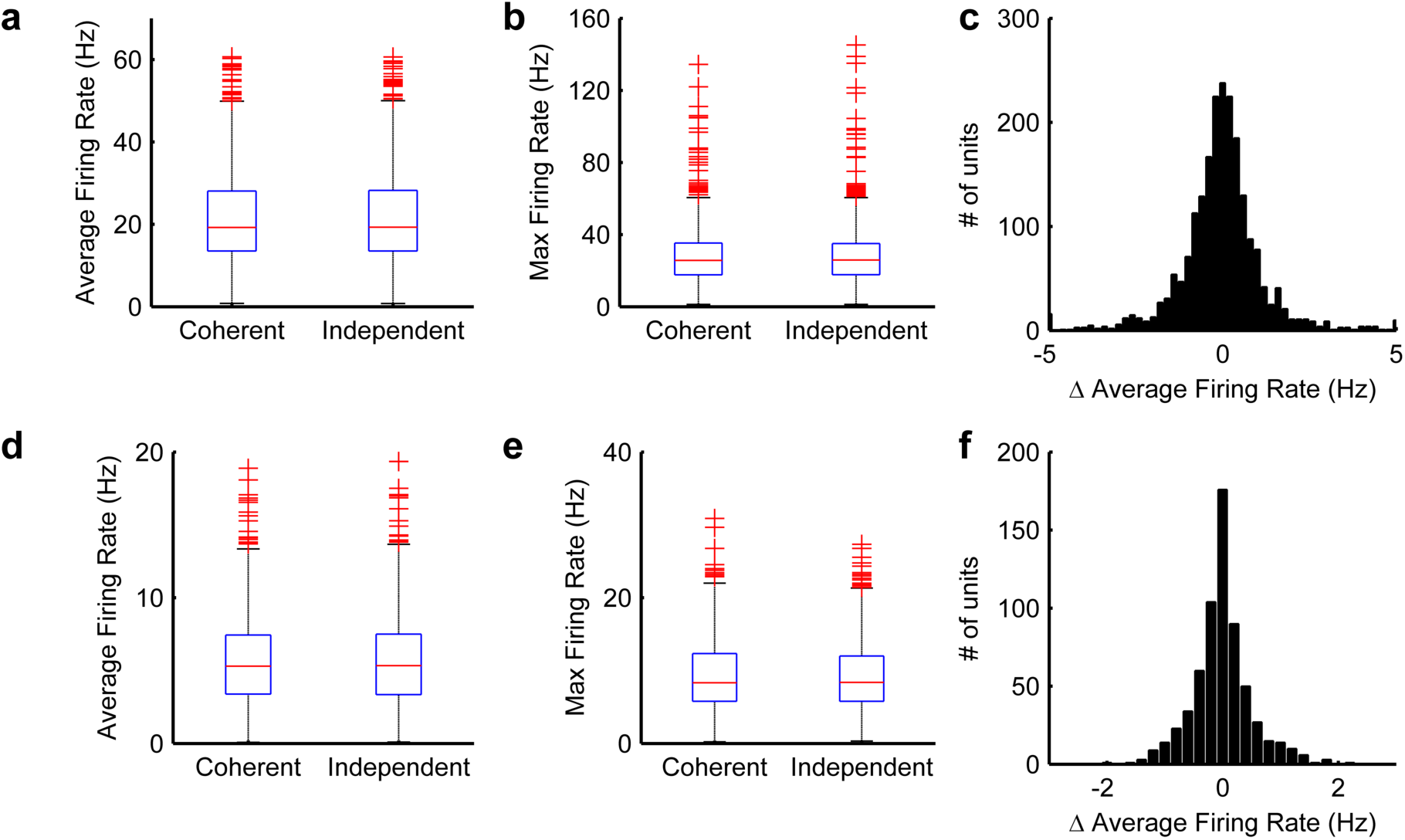
There were no statistically significant changes in mean **(a,d)** or max **(b,e**) firing rate between temporally coherent and temporally independent datasets in either the awake **(a,b,c)** or anesthetised **(d,e,f)** datasets**. Awake dataset:** For all units: Mean firing rate t_540_= -0.0308, p =0.9754; Max firing rate: t_540_ = 0.4354, p =0.6636. For units with a significant VPI value: Mean: t_180_= -0.0631, p =0.7694; Max: t_180_= 0.8563, p =0.0939. **Anesthetised dataset:** all units: Mean firing rate t_-664_ = -0.0638, p = 0.9492. Max firing rate: t_664_= 0.0047, p =0.9947. Significant VPI units Mean firing rate t_348_ 0.0308, p =0.9498. Max firing rate: t_348_= 0.0235, p =0.9912.

**Supplemental Figure 6 (related to Figure 4).**
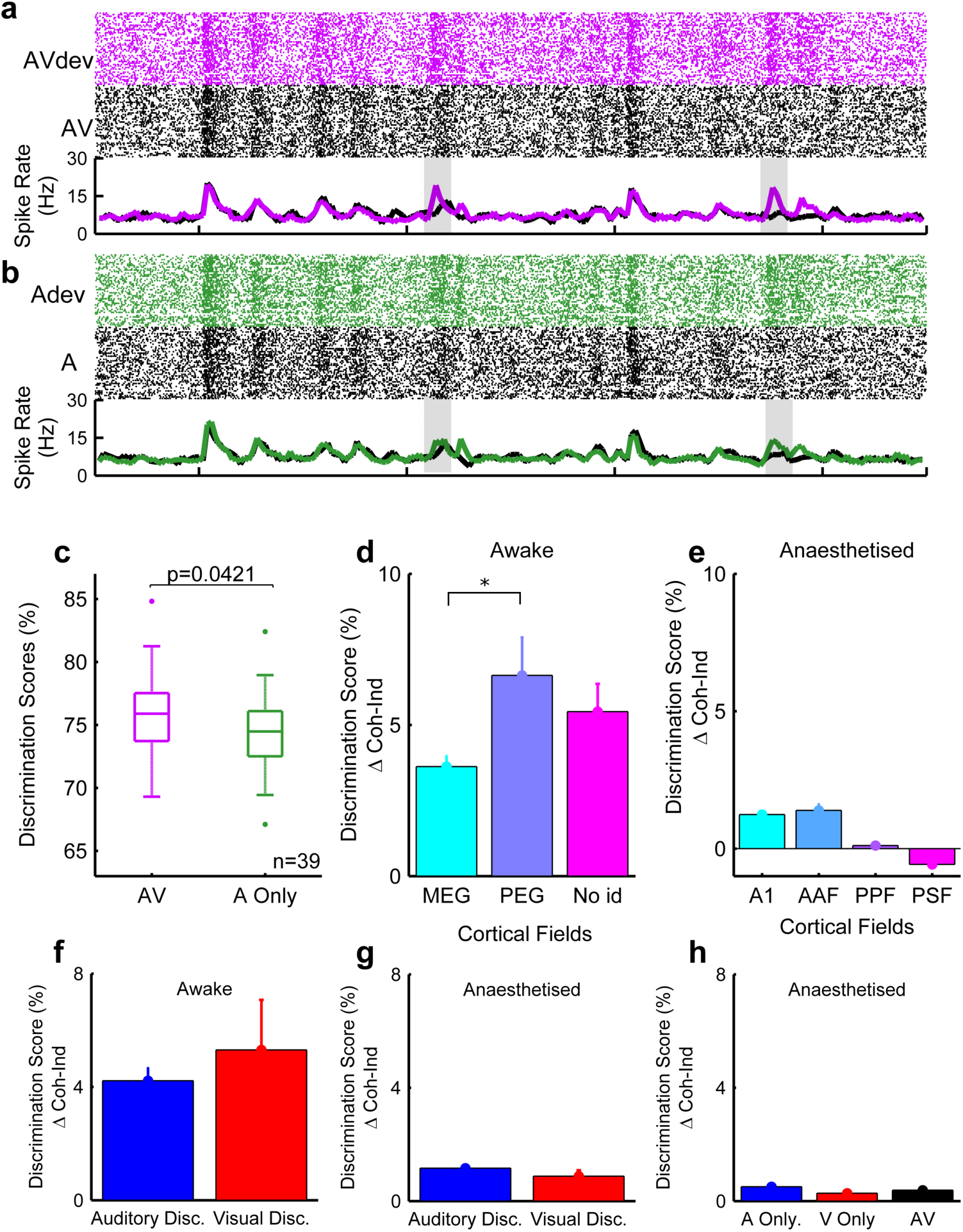
**a,b**, Rasters and PSTH for single stream stimuli containing deviants (top row, black) or without deviants (bottom row, purple/green). **a**, auditory stream presented with a temporally coherent visual stimulus, **b**, auditory stream presented in isolation. Grey panels indicate the timing of the timbre deviants. **c**, discrimination scores for detecting trials with deviants in them were significantly higher in AV trials than A only trials. Recordings were made in awake animals (# of animals =3, n=39 driven units,).Pairwise comparison t_76_ = 2.0676, p = 0.0421) **d,e,** comparison of how visual temporal coherence influenced deviant encoding across cortical fields in awake data and anesthetised data. In awake data, a one-way ANOVA across cortical fields showed a significant effect of field (F (2,165) = 2.6710, p= 0.0322) with post-hoc comparisons indicating that the discrimination scores were greater in the PEG than MEG. However, there were no significant effect of field in anaesthetised data (F (3,245) = 2.0627, p = 0.1057) f,g No differences were found in discrimination score between auditory discriminating and visual discriminating units in awake recordings (F (1,100) = 0.2547, p = 0.6149) or in anaesthetised recordings (F (1,119) = 0.0689, p = 0.7933). h There were also no differences across different unit type in anaesthetised recording (F (2,423) = 0.846, p = 0.9169)

**Supplemental Figure 7 (related to Figure 6).**
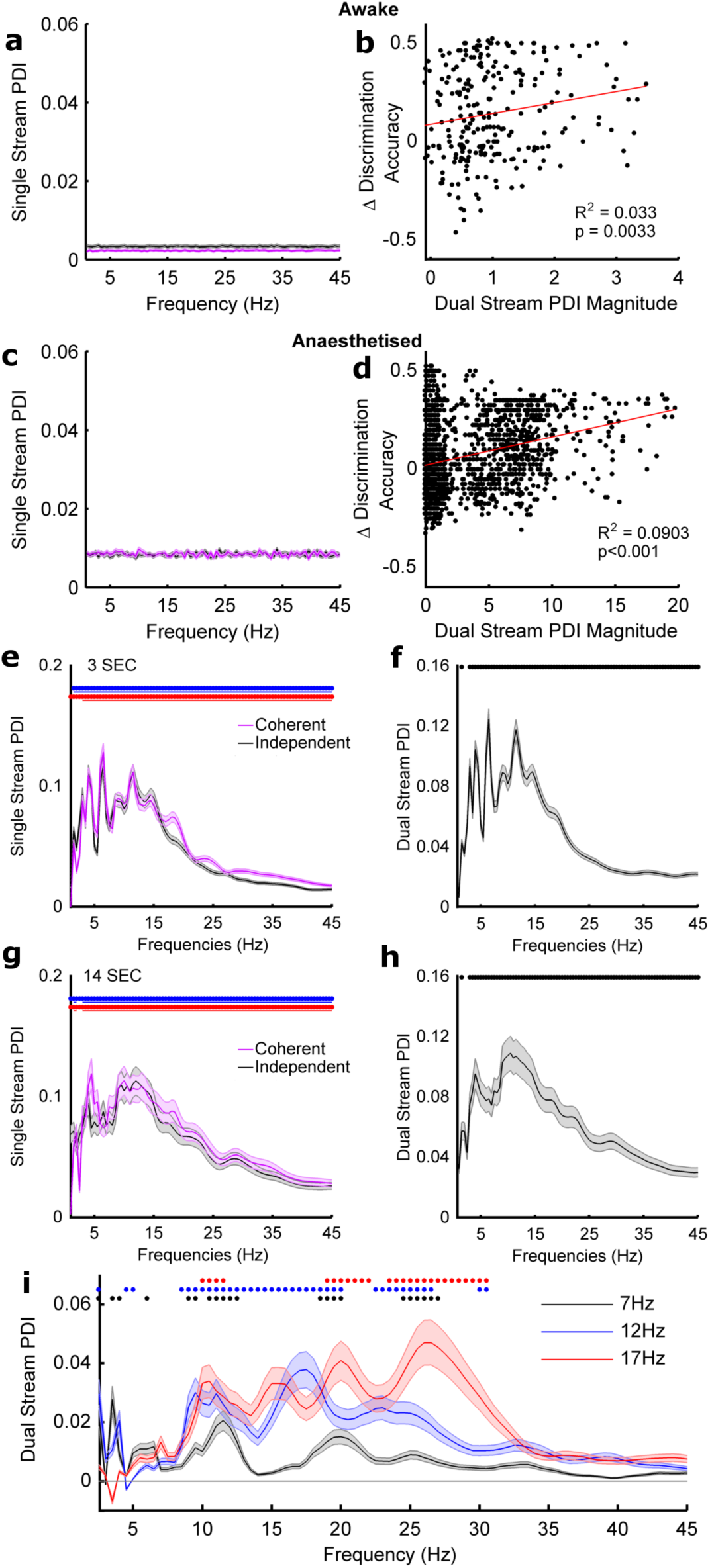
**a,c** dual stream power discriminability index values for awake (a) and anesthetised datasets (c). In neither case was there any frequency whose power was significantly influenced by the visual stimulus identity. **b,d**, relationship between phase discriminability index (PDI) values measured in the dual stream condition and the VPI score. There is a weak correlation between the magnitude of the dual stream PDI values and VPI values. During our initial analysis we observed that PDI values were higher in anesthetised animals than awake animals. In order to determine whether this was a difference due to behavioural state or simply an artefact of stimulus length for all of the analysis reported in this paper we restricted analysis of the anesthetised responses to the first three seconds of the stimulus. In **e-h** we explicitly compare the PDI values obtained for 3 second (**e,f**) and 14 second (**g,h**) single stream (**e,g**) and dual stream (**f,h**) stimuli. to match that recorded in the awake dataset. While phase coherence values were slightly higher for longer duration stimuli and hence at longer stimulus durations the ITPC profile and resulting PDI varied more smoothly with frequency. However at both durations phase values were significantly different from zero at all frequencies. The pattern of significant phase selectivity values was also preserved across stimulus durations. (**b, d**). Frequency points at which the single stream PDI value and dual stream PDI values were similar in 3 second length (**a, b**) and 14 second length (**c, d**) Blue, red and black symbols indicate where the PDI was significant (pairwise t-test, a = 0.0012 with bonferoni correction). **i** Dual stream stimuli were generated with three different amplitude modulation rates (<7Hz, as in the main experiment, <12Hz and <17Hz, values picked to avoid harmonics of 7 Hz) and responses to these were recorded in 92 units. Symbols indicate where the dual stream phase selectivity index was significant (pairwise t-test, p < 0.05 with correction). In all three cases significant phase coherence is seen between 10Hz-11.5Hz, 19Hz-20Hz and 24-26 Hz.

